# Mosquitoes (Diptera: Culicidae) of Ecuador: A revised checklist, new records and species of medical importance

**DOI:** 10.1101/2021.02.10.429771

**Authors:** Patricio Ponce, Varsovia Cevallos, Andrés Carrazco-Montalvo

## Abstract

Ecuador is one of the countries with the highest biodiversity in the world, this can also be seen in the Culicidae family. However, there are a limited number of studies on this group. This work provides a baseline reference for this insect group and information about numerous potential and vector disease species. Species names and records were extracted from the National Mosquito Reference Collection at INSPI-Quito, published literature and web databases. The specimens at the INSPI collection were identified using morphological keys and, in a few cases, using molecular markers in the genus Anopheles. An updated list includes the subfamilies Culicinae and Anophelinae, eight tribes, 22 genera and 200 species. We present 18 species cataloged as new records for Ecuador represented in two subfamilies, 6 tribes and 9 genera. Taxonomic notes, geographical distribution and medical importance data for the species involved are provided. The updated list of Mosquitoes (Diptera: Culicidae) is a guide for researchers and health personnel when studying biodiversity, fauna, insect vectors and strategies to prevent the spread of vector diseases.

## Introduction

Ecuador is considered one of the most biodiverse countries in the world due to rich and unique habitats. Arthropods dominate rainforest ecosystems in terms of species numbers and Ecuador is among the countries recognized as one of the centers of speciation, because of its complex geological history and climate (1).

Ecuador is located on the northwest coast of South America, and its territory includes the continental mainland and the Galápagos Islands in the Pacific Ocean. The mainland of Ecuador is divided geographically into the coastal lowlands on the west of the Andes Mountains, the Andes or highlands, with valleys and mountains with snowed peaks of up to 6300 m, and lowlands on the east of the Andes that are part of the Amazon basin. As a consequence of this geographical diversity Ecuador has 26 Holdridge ecological life zones, one of the highest in the Neotropics (2, 3). These characteristics make Ecuador one of the most biodiverse countries in the world. This biological richness is apparent also in the Culicidae family. However, in Ecuador, this group has been overlooked for the last 50 years. Just recently, records of new species have been reported for the country (4–6).

There are around 3570 species of the family Culicidae (Diptera) in the world, distributed in 112 genera and 145 subgenera, and arranged in two subfamilies, Anophelinae and Culicinae. Anophelinae has three genera and Culicinae 109 genera (7). The first descriptions of mosquitoes from Ecuador were done by Alexander von Humbolt in 1819 (8). The study of the mosquito fauna of Ecuador then continued by Francisco Campos who contributed several publications on mosquitoes (9–12). Campos collected mosquitoes mainly around the city of Guayaquil and coastal areas. Those specimens were sent to Harrison G. Dyar, who described several species from Ecuador around 1918-1925 (13) and several species were named after Campos.

Levi-Castillo has been, so far, one of the most significant contributors to the knowledge of mosquitos of Ecuador. Among his most important publications are the Atlas de los Anofelinos Sudamericanos (14) and the first Culicidae checklist of the country (15). In 1962, Levi-Castillo abruptly quit entomology, which resulted in almost no publications dealing with the mosquito fauna from Ecuador since then. In the next decade, 28 species of Culicidae for Ecuador were reported (8), 19 of which were described by Levi-Castillo. The latter author reported an extended list of mosquitoes collected mainly on the coastal areas and the highlands (15). Heinemann and Belkin (16) reported detailed data for 80 mosquito species from Ecuador. The last list reported for Ecuador mentions 179 species (4).

This work provides a baseline reference for this insect group and information about numerous potential and vector disease species.

## Materials and Methods

Species names and records were extracted from the National Mosquito Reference Collection (CIREV) at INSPI-Quito (Instituto Nacional de Investigación en Salud Publica), here cited as INSPI collection, published literature and web databases (17–19). The specimens at the INSPI collection were identified using morphological keys and, in a few cases, using molecular markers in the genus *Anopheles.* The information was cross-checked to include only valid species names that are verified in primary bibliographic reports. Most of the species records by Levi-Castillo were not included, since most of the specimens used to describe or report the species are missing. The names used in this paper are the accepted names published by Harbach and Kitching (20) and genera and subgenera abbreviations according to Reinert (21, 22).

The species are presented in alphabetical order (Appendix 1) with synonym names and the geographic distribution in Ecuador. A list of new records is also presented (Table 1). Highland mosquito species denomination is arbitrary when insects were collected from over 1000 m of altitude. The medical importance of the pertinent species is also mentioned. The list treats the taxa *Lutzia* and *Onirion* as genera, although there is still discussion regarding their taxonomic status (23, 24).

**Table 1.**
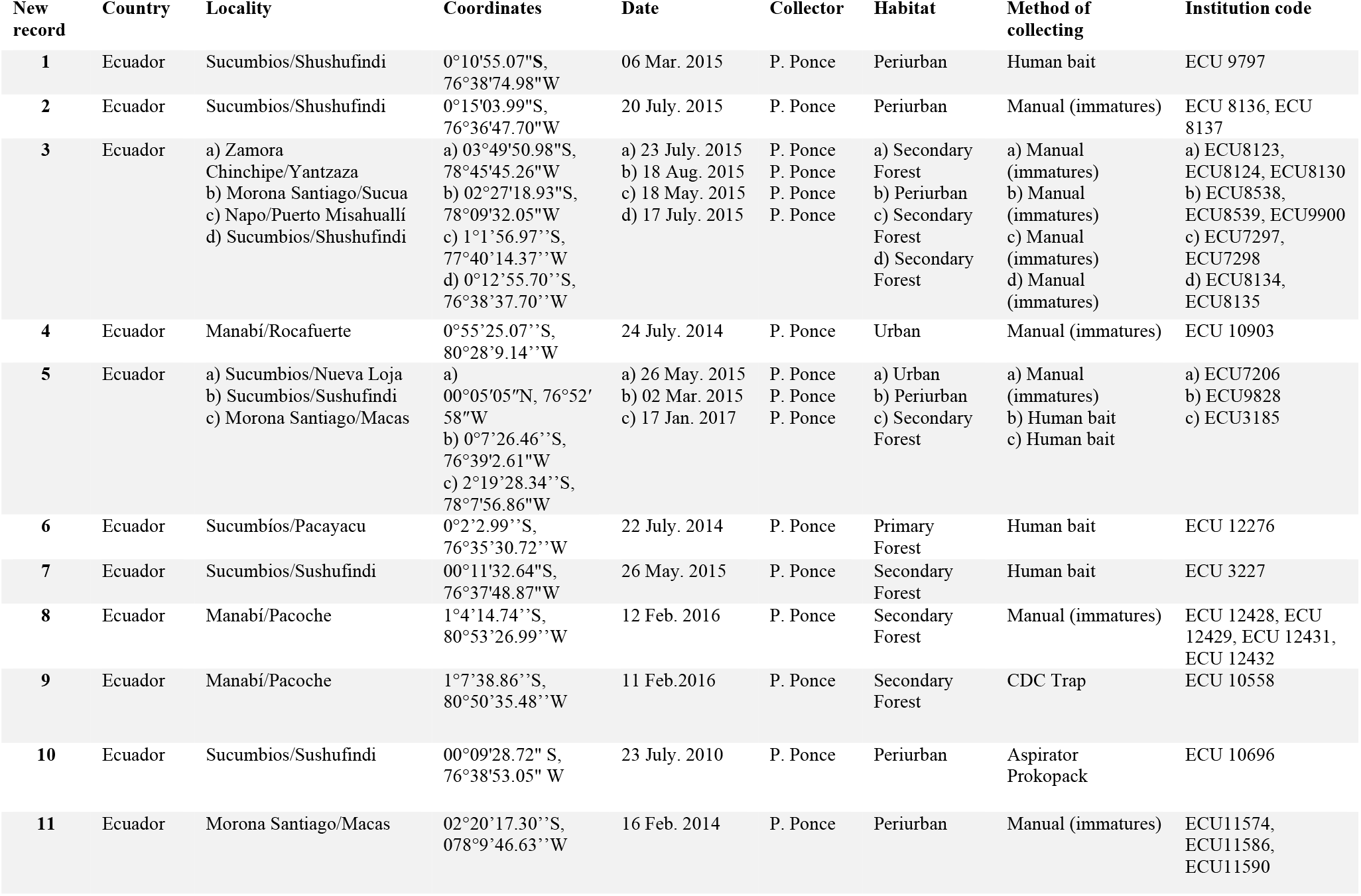
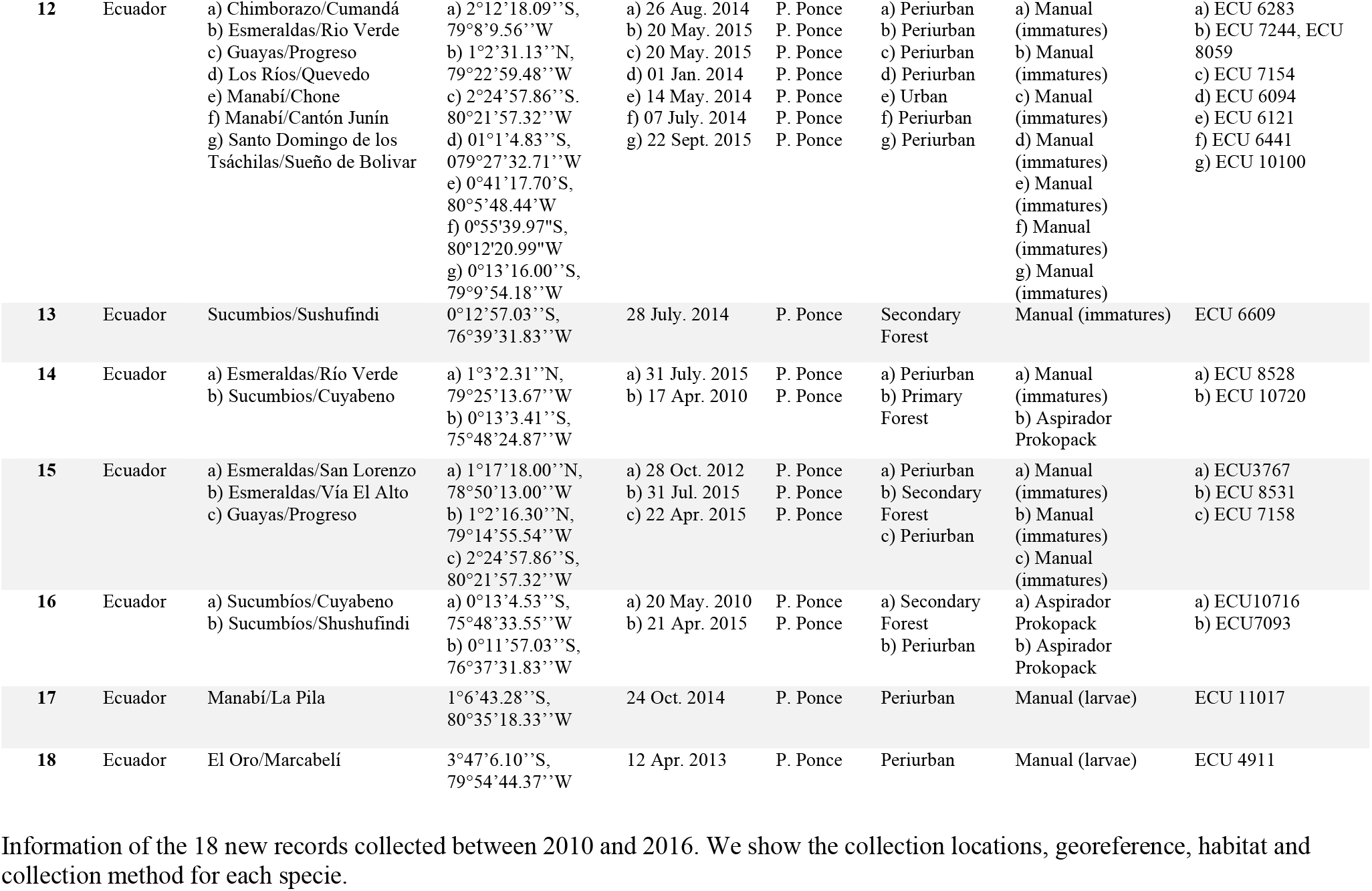
Information of New Records

## Results

The updated list presents 200 species in 22 genera of Culicidae with geographical distribution and georeferenced points (Appendix 1). The subfamily Culicinae is represented by 8 tribes: Aedeomyiini (*Aedeomyia*), Aedini (*Aedes, Haemagogus* and *Psorophora*), Culicini (*Culex, Deinocerites, Galindomyia* and *Lutzia*), Mansoniini (*Coquillettidia, Mansonia*), Orthopodomyiini (*Orthopodomyia*), Sabethini (*Johnbelkinia, Limatus, Onirion, Runchomyia, Sabethes, Trichoprosopon* and *Wyeomyia*), Toxorhynchitini (*Toxorhynchites*), and Uranotaeniini (*Uranotaenia*). The Anophelinae subfamily is represented by the genera *Anopheles* and *Chagasia*.

We report 24 species of *Anopheles,* three of *Chagasia,* one of *Aedeomyia,* 21 of *Aedes,* 11 of *Haemagogus*, 12 of *Psorophora*, 61 of *Culex*, one of *Deinocerites*, one of the genus *Galindomyia*, one of *Lutzia*, five of *Coquillettidia*, five of *Mansonia*, three of *Orthopodomyia*, two of *Johnbelkinia*, three of *Limatus*, one of *Onirion*, one of *Runchomyia*, 14 of *Sabethes*, six of *Trichoprosopon*, 14 of *Wyeomyia*, five of *Toxorhynchites* and five of *Uranotaenia*.

We present below 18 species cataloged as new records for Ecuador represented in two subfamilies, 6 tribes and 9 genera. These mosquitoes were collected between 2010 and 2016. For these species, the collection sites, geographic coordinates, collection dates, habitat and collection method are detailed (Table 1). The geographical distribution is in several localities of the country and is represented by a map (Figure 1).

**Figure 1.**
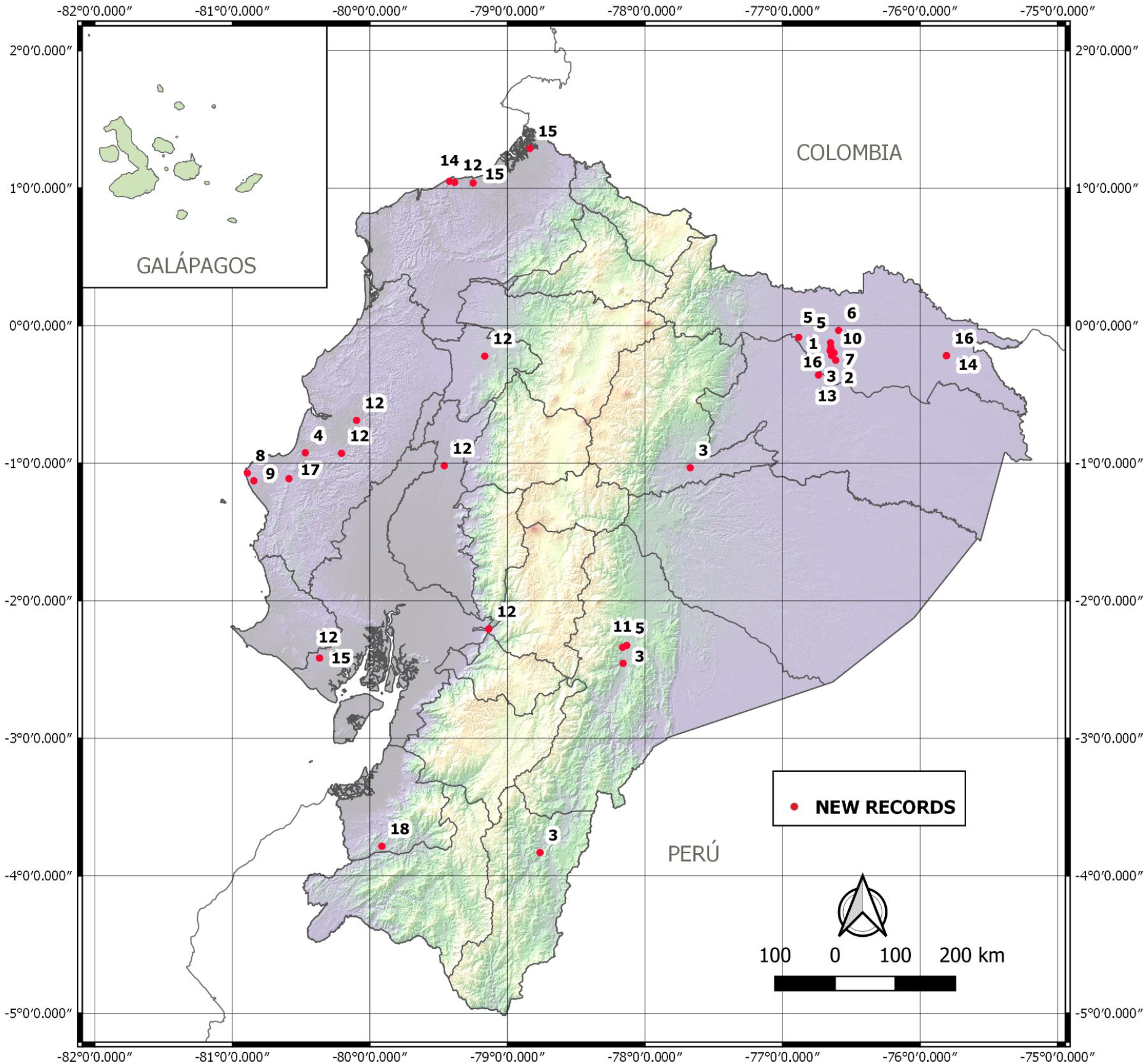
Geographical distribution of the new records in Ecuador. Each individual is represented with the New Records according to Table 1.

### Subfamily Anophelinae

- 1. *Anopheles (Nyssorhynchus) darlingi* Root, 1926. Synonym*: Anopheles paulistensis* Galvao, Lane & Correa, 1937. (1♀);
- 2. *Anopheles (Nyssorhynchus) evansae* Brèthes, 1926. Synonym: *Anopheles noroestensis* Galvao & Lane, 1937. (2♀);
- 3. *Anopheles (Nyssorhynchus) nuneztovari* Gabaldon, 1940. Synonym: *Anopheles goeldii* Rozeboom & Gabaldon, 1941. (7♀, 3♂).

### Subfamily Culicinae. Tribe Aedini

- 4. *Aedes (Ochlerotatus) albifasciatus* (Macquart, 1838). Synonyms: *Aedes colonarius* Dyar, 1924; *Culex albifasciatus* Macquart, 1838; *Culex annuliferus* Blanchard, 1852; *Culex flavipes* Macquart, 1838. (1♂);
- 5. *Psorophora (Grabhamia) dimidiata* Cerqueira, 1943. (3♀);
- 6. *Psorophora (Psorophora) holmbergii* Lynch Arribalzaga, 1891. Synonym: *Psorophora agoggylia* Dyar, 1922. (1♀).

### Tribe Culicini

- 7. *Culex (Aedinus) accelerans* Root, 1927. Synonym: *Culex paraplesia* Dyar, 1922. (1♀);
- 8. *Culex (Culex) stigmatosoma* Dyar, 1907. (3♂, 1♀);
- 9. *Culex (Culex) thriambus* Dyar, 1921. Synonyms: *Culex affinis* Adams, 1903; *Culex peus* Speiser, 1904; *Culex eumimetes* Dyar & Knab, 1908. (1♀).

### Tribe Mansoniini

- 10. *Coquillettidia (Rhynchotaenia) albicosta* (Peryassu, 1908). Synonym: *Taeniorhynchus albicosta* Peryassu, 1908. (1♀).

### Tribe Orthopodomyiini

- 11. *Orthopodomyia albicosta* (Lutz, 1904). Synonym: *Bancroftia albicosta* Lutz, 1904. (3♀).

### Tribe Sabethini

- 12. *Sabethes (Sabethes) albiprivus* Theobald, 1903. Synonyms: *Sabethes albiprivatus* Lutz, 1904; *Sabethes albiprivatus* Theobald, 1907; *Sabethes neivai Petrocchi,* 1927. (6♀, 2♂);
- 13. *Sabethes (Sabethes) belisarioi* Neiva, 1908. Synonyms: *Sabethes argyronotum* Edwards, 1928; *Sabethes goeldii* Howard, Dyar & Knab, 1917; *Sabethes schausi* Dyar & Knab, 1908. (1♀);
- 14. *Sabethes (Sabethes) purpureus* (Theobald, 1901). Synonyms: *Sabethes purpureus* Peryassu, 1908; *Sabethes remipusculus* Dyar, 1924; *Sabethoides purpureus* Theobald, 1907. (1♀, 1♂);
- 15. *Sabethes (Sabethes) quasicyaneus* Peryassú, 1922. (2♀, 1♂);
- 16. *Sabethes (Sabethoides) glaucodaemon* Dyar & Shannon, 1925. Synonym: *Sabethoides glaucodaemon* Dyar & Shannon, 1925. (2♀).

### Tribe Toxorhynchitini

- 17. *Toxorhynchites (Lynchiella) guadeloupensis* (Dyar & Knab, 1906). Synonyms: *Megarrhina horei* Gordon & Evans, 1922; *Megarhinus tucumanus* Brèthes, 1926; *Megarhinus arborealis* Shannon & Ponte, 1927; *Megarrhina guianensis* Bonne-Wepster & Bonne, 1953. (1♂);
- 18. *Uranotaenia* (*Uranotaenia*) *ditaenionota* Prado, 1931. Synonym: *Uranotaenia burkii* Lane, 1936. (1♀).

### Species of medical importance

Subfamily Anophelinae Grassi, 1900.

The number of species of the genus *Anopheles* is the second largest in Ecuador. Out of the 472 formally recognized species of this subfamily (7, 25), 24 species (5%) have been recorded in Ecuador. In comparison, Colombia has 48 species (26, 27), and Peru 43 (28).

In Ecuador, there are 17 species of *Anopheles* vectors and potential vectors for malaria out of the 34 species of *Anopheles* that are reported in the Americas (28–34). The species present in Ecuador that have been reported as infected with the malaria parasites in the Americas are: *Anopheles (Anopheles) calderoni* Wilkerson, 1991; *Anopheles (Anopheles) fluminensis* Root, 1927; *Anopheles (Anopheles) mattogrossensis* Lutz & Neiva, 1911; *Anopheles (Anopheles) neomaculipalpus* Curry, 1931; *Anopheles (Anopheles) pseudopunctipennis* Theobald, 1901; *Anopheles (Anopheles) punctimacula* Dyar & Knab, 1906; *Anopheles (Kerteszia) lepidotus* Zavortink, 1973; *Anopheles (Kerteszia) neivai* Howard, Dyar & Knab, 1913; *Anopheles (Nyssorhynchus) albimanus* Wiedemann, 1820; *Anopheles (Nyssorhynchus) aquasalis* Curry, 1932; *Anopheles (Nyssorhynchus) darlingi* Root, 1926; *Anopheles (Nyssorhynchus) evansae* Brèthes, 1926; *Anopheles (Nyssorhynchus) nuneztovari s.l.* Gabaldon, 1940; *Anopheles (Nyssorhynchus) oswaldoi* (Peryassú, 1922); *Anopheles (Nyssorhynchus) rangeli* Gabaldón, Cova García & Lopez, 1940; *Anopheles* (*Nyssorhynchus*) *triannulatus* (Neiva & Pinto, 1922); *Anopheles (Nyssorhynchus) trinkae* Faran, 1979.

Subfamily Culicinae Meigen, 1818.

*Aedeomyia (Aedomyia) squamipennis* (Lynch Arribálzaga, 1878) is the only species of this genus in Ecuador, and it has been recorded only on the west lowlands of the Andes (35, 36). This species has been incriminated as carrier of 24 strains of virus related serologically to prototype Gamboa in Ecuador (37, 38). This species has also been identified as vector of avian malaria (39) (39).

There are 21 species of *Aedes* registered for Ecuador. *Aedes aegypti* (Linnaeus, 1762) has been reported from continental eastern and western sides of the Andes, and the Galapagos Islands. This species has been reported from localities of up to 1650 m over sea level. This species is known for the dispersal of viruses including Chikungunya (CHKV), Dengue (DENV) and Zika (ZIKV) among the human population (40). *Aedes aegypti* has also shown midgut infection by Sindbis virus (41, 42). This species is also a suspected as vector of *Wuchereria bancrofti* (Raghavan, 1961).

*Aedes albopictus* was recorded in Ecuador in late 2017 in Guayaquil and has already spread to the Amazon basin and northwestern lowlands (6). This species has been incriminated as a vector of several viruses Chikungunya (CHKV), Dengue (DENV), and as a potential vector of Zika (ZIKV), Eastern equine encephalitis (EEEV), La Crosse (LACV), Venezuelan equine encephalitis (VEEV), Japanese encephalitis (JEV), St. Louis encephalitis, West Nile (WNV), and Yellow fever (YFV), and as a vector of filarial parasites (43).

*Aedes taeniorhynchus* (Wiedemann, 1821) is vector of Venezuelan equine encephalitis virus (VEEV) (44, 45). This species is considered a potential threat for the Galapagos Islands fauna as a competent vector of West Nile virus (WNV) (46).

There are eleven species of *Haemagogus* mosquitoes reported for Ecuador. *Haemagogus leucocelaenus* (Dyar & Shannon, 1924) is a vector of yellow fever virus (47, 48). *Haemagogus anastasionis* Dyar, 1921 has been reported for Ecuador and it is a potential vector of yellow fever and Mayaro virus (5).

Thirteen species of the genus *Psorophora* have been reported for Ecuador. *Psorophora ferox* (Humboldt, 1819) has been reported as carrier of Ilheus virus (ILHV) in Central, South America and Trinidad and Tobago. In Ecuador the virus isolate was obtained from a human, however, this report needs to be confirmed due to cross-reactivity in serologic assays to other flaviviruses (49).

The genus *Culex* has 61 species in Ecuador, representing 8% of the world diversity. There are 14 species (19%) exclusively in the western lowlands of Ecuador, while 31 species (43%), of the species are in the eastern lowlands, 13 species (18%) distributed on both sides of the Andean mountains, and three species (4%) are in the highlands.

*Culex (Culex) quinquefasciatus* Say, 1823. This species is widely distributed on the west and east of the Andean mountains of Ecuador, and has been introduced into the Galapagos Islands (15, 50). It is the most important species for transmitting filariasis and Oropouche virus (51), *Culex quinquefasciatus* is an established species in the Galapagos Islands and it has been reported as vector of avian malaria and avian pox virus. It is also a competent vector of the West Nile Virus (50).

*Culex nigripalpus* Theobald, 1901. This mosquito has been reported as responsible of West Nile Virus transmission to chickens (52).

*Culex iolambdis* Dyar, 1918 is a carrier of Venezuela encephalitis virus, however, it is unknown if it is able to transmit it (53).

*Culex (Melanoconion) ocossa* Dyar & Knab, 1919 carries a flavivirus in Peru (54) and the Venezuelan equine encephalitis virus complex (55).

The genus Deinocerites, is represented by one specie reported in Ecuador*. Deinocerites pseudes* Dyar & Knab, 1909 apparently is a vector of Venezuelan and St. Louis equine encephalitis (56).

Tribe Mansoniini

Mosquitoes of the genus *Coquillettidia* have been reported as carriers of avian malaria parasites (57) and several viruses including Eastern equine encephalomyelitis virus (EEE), West Nile virus (WNV), Usutu virus (USUV), Tahyna virus (TAHV), Sindbis virus (SINV), Batai virus (BATV) (58). *Coquillettidia albicosta* present in Ecuador was reported as carrier of *Rochambeau* virus (RBUV) in French Guiana in 1973 (59).

The genus *Mansonia* Blanchard, 1901 is represented by three species reported in Ecuador. *Mansonia titillans* (Walker, 1948) is known to carry the virus of Venezuelan equine encephalitis (55) and it may also transport eggs of *Dermatobia hominis* (60–63).

The genus *Johnbelkinia* Zavortink, 1979 is represented by two species. *Johnbelkinia ulopus* has been mentioned as a potential vector of arbovirus (64, 65) and is known to carry eggs of *Dermatobia hominis* in Colombia (66).

There are three species of the genus *Limatus* Theobald, 1901. *Limatus durhami* and *Li. flavisetosus* have been reported carrying viruses (63).

There are 14 species of *Sabethes* reported for Ecuador. *Sabethes chloropterus* has been reported as vector of yellow fever (67) and other arboviruses like St. Louis encephalitis and Ilhéus (60, 63). *Sabethes amazonicus* has been indicated as a potential vector of yellow fever virus (5).

There are 14 species of the genus *Wyeomyia* Theobald, 1901 from Ecuador. There are reports of virus presence in species of this genus, however, it is not known their medical importance (60, 63).

## Discussion

The updated mosquito species present in Ecuador, along with the specimens deposited in a National Reference Collection, will help to understand the diversity and the dynamics of several diseases transmitted by species of this group. Presently, the National Mosquito Reference Collection (CIREV) at INSPI - Quito holds 98 species with detailed collection data, out of the 200 reported for Ecuador. The list presented, most likely, is a partial inventory of the Ecuadorian mosquito fauna, considering that the country has one of the highest biodiversity in the region (68) and many ecosystems have not been fully surveyed. Additionally, the updated record of mosquito species in continental Ecuador is also relevant for the surveillance of invasive species in fragile ecosystems like the Galapagos Islands. Extensive collections are further needed and taxonomic and phylogenetic tools should be applied to establish the distribution, identity and disease incrimination of species complexes.

The number of mosquito species for Ecuador is significant considering the country size, for instance Brazil has 447 species of mosquitoes, while Mexico has about 225, Colombia 324 and Peru 129 species (27, 69). Foley et al. reported only 118 species, 23 endemic species and 34 types for Ecuador (69). The most recent list of Culicidae of Ecuador (4) reports 179 species for Ecuador. The number of mosquito species reported in this revision represents 6% of the Culicidae species. Out of the 200 species, 13 (6.5%) have Ecuador as the type locality. Mosquito taxonomy studies are still scarce in Ecuador and there are not published morphological identification keys available for local mosquito fauna. The taxonomic status of several species presents in Ecuador, in particular of the genus *Anopheles* that has several species complexes, needs to be revised along with their vector disease status. According to the World Health Organization *Anopheles albimanus, An. punctimacula* and *An. pseudopunctipennis* are the main malaria vectors in Ecuador (70). However, there are no reports of *Anopheles* species infected with the protozoan parasite in Ecuador.

Comprehensive data about the mosquito fauna and its distribution in Ecuador will help vector control officials to prevent the spread of diseases.

## Conclusion

The updated list of Mosquitoes (Diptera: Culicidae) present in Ecuador, with the new records and georeferencing; provides information on group diversity, geographic distribution, and medical importance of some species. This is a guide for researchers and health personnel when studying biodiversity, fauna, insect vectors and strategies to prevent the spread of vector diseases.

## Acknowledgements

We thank Giovanni Onore and Stephany Villota for revising the manuscript. We thank Carlos Yánez for formatting the coordinates. This research was partially funded by Senescyt, grant PIC-12-INH-003, Agreement 20120468.

## Appendix 1

List of valid culicid species reported for Ecuador. The sub-generic classification, author, year of publication, localities where recorded and literature references for each species are included. Species distribution in the East (E), West (W), highlands (H) and the Galápagos Islands (G) with the collection reference numbers (ECU). NR indicates new record for Ecuador. ** indicates endemic species. † indicates species with type locality in Ecuador.

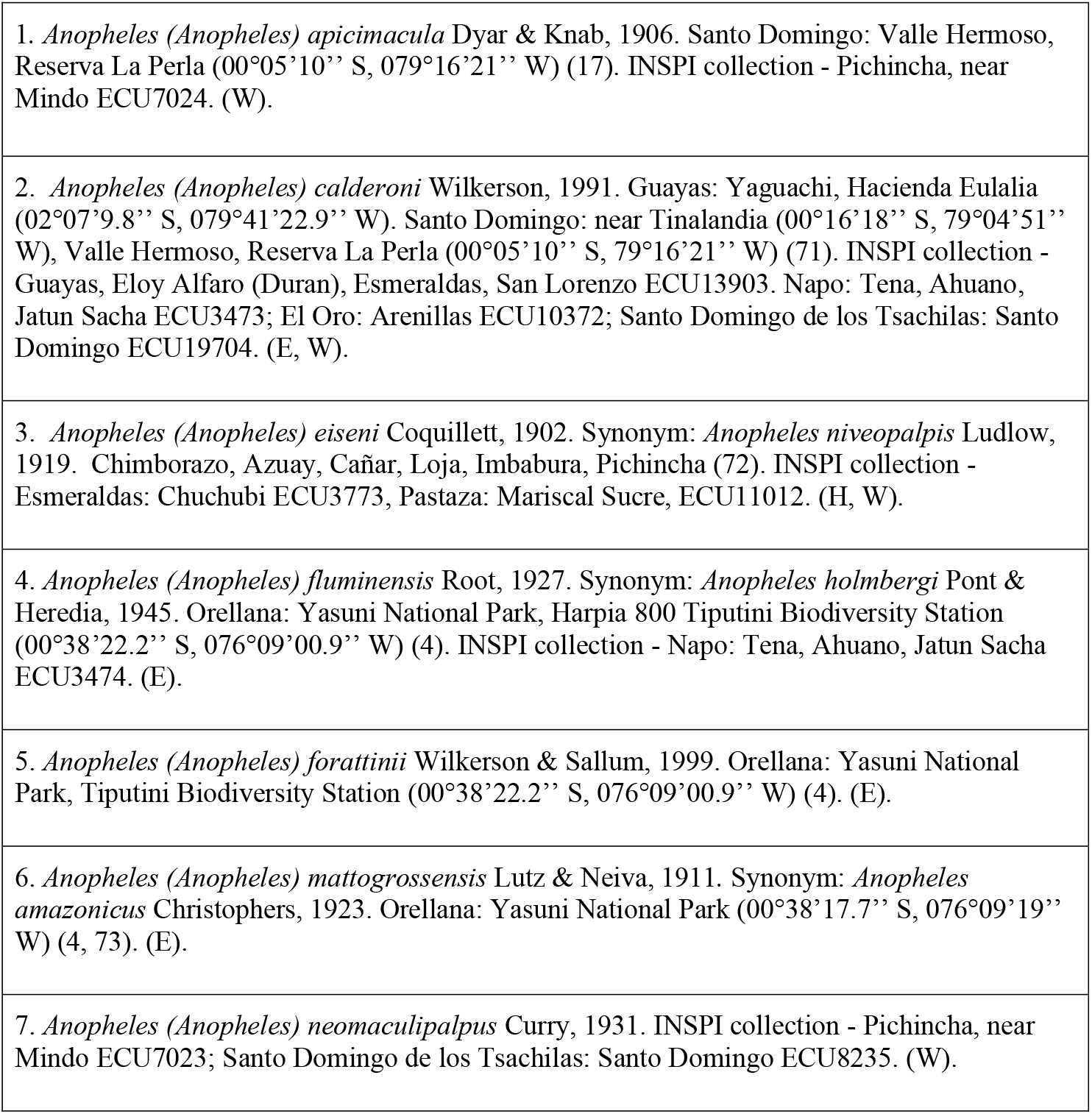
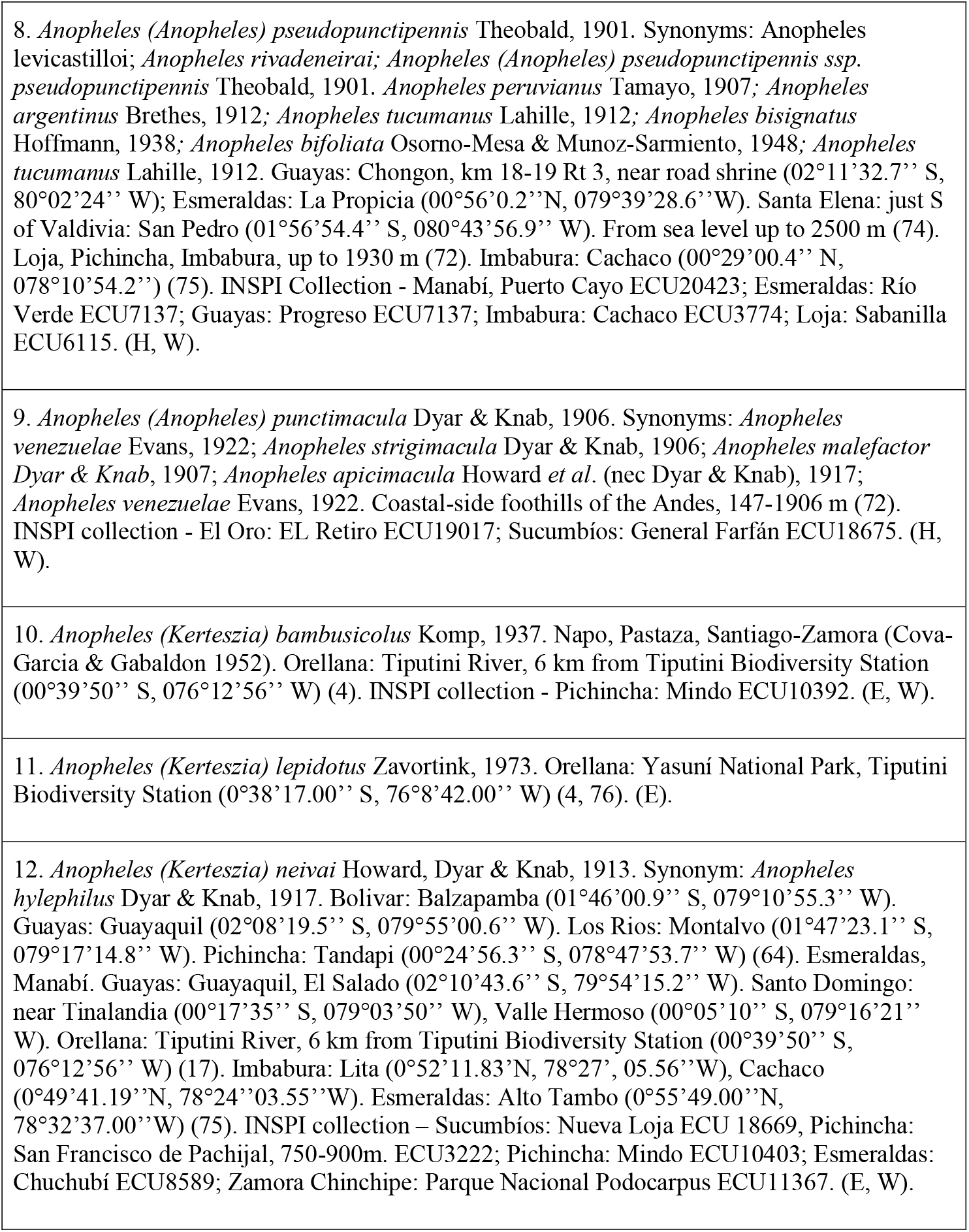
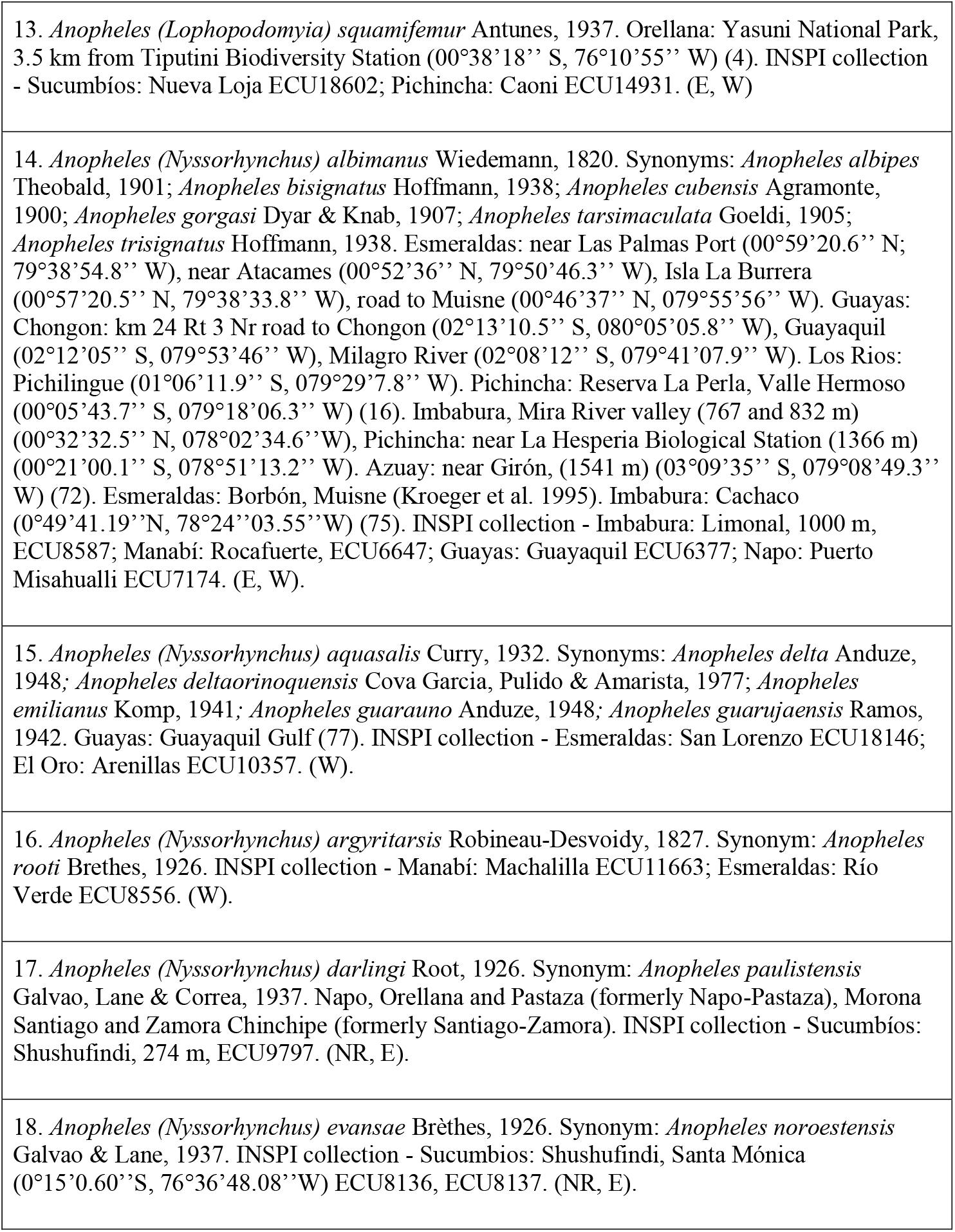
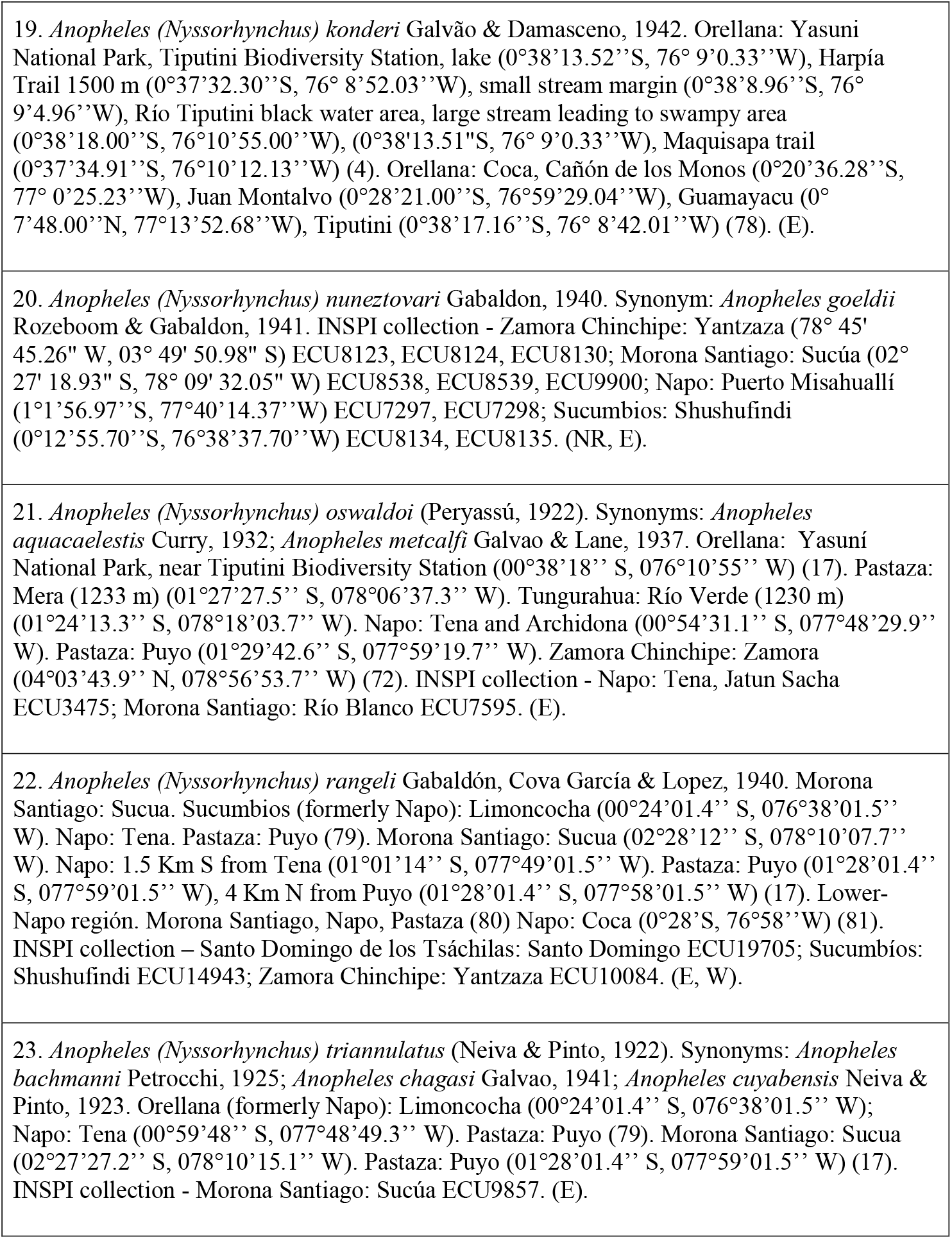
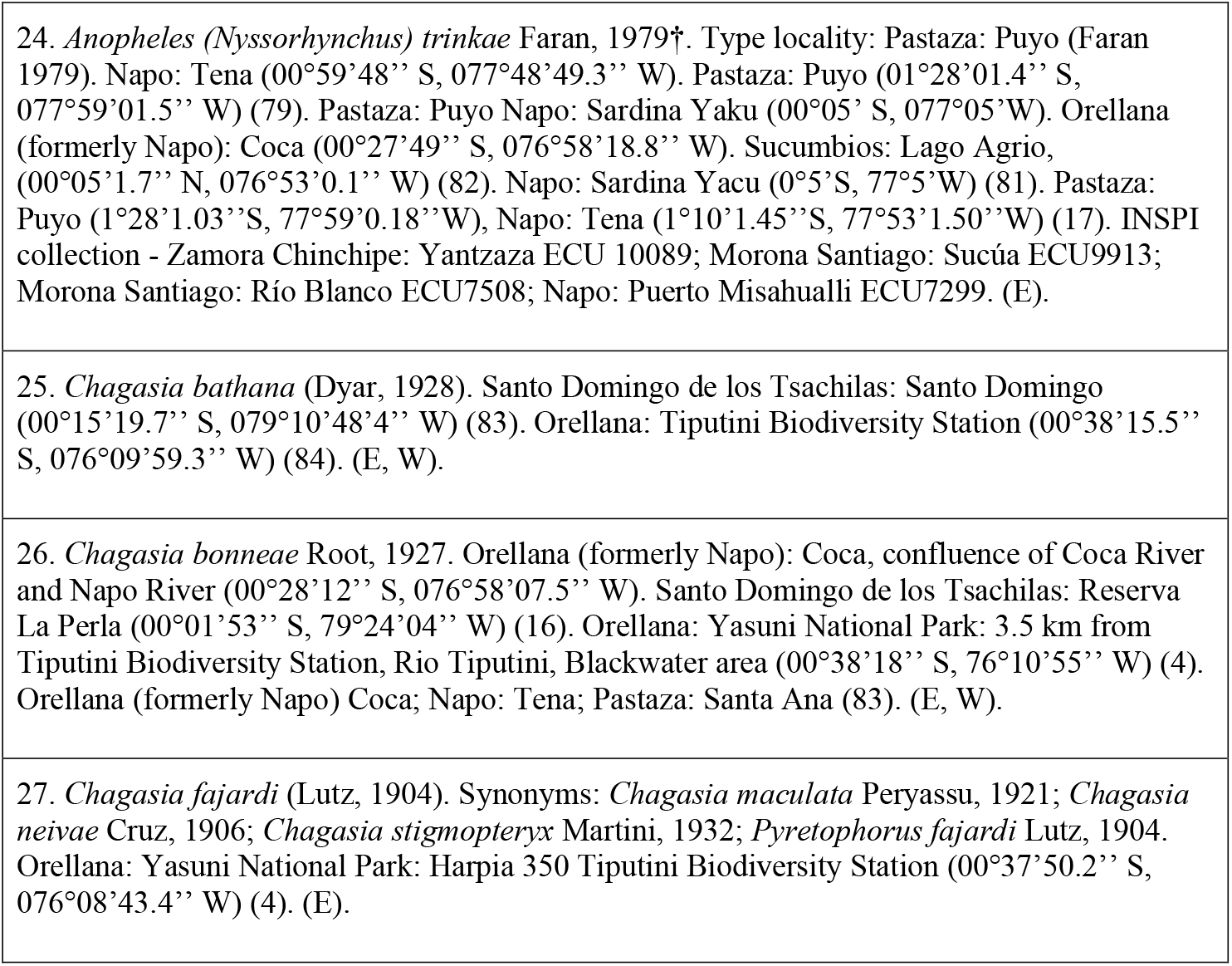
- Subfamily Anophelinae: 27 species

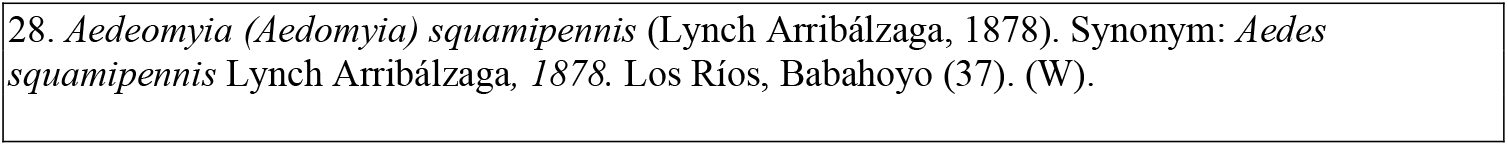
- Subfamily Culicinae. Tribe Aedeomyiini: 1 specie

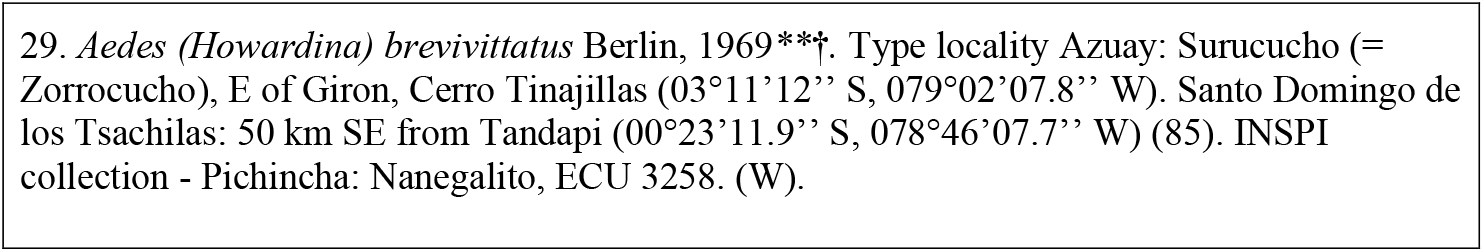
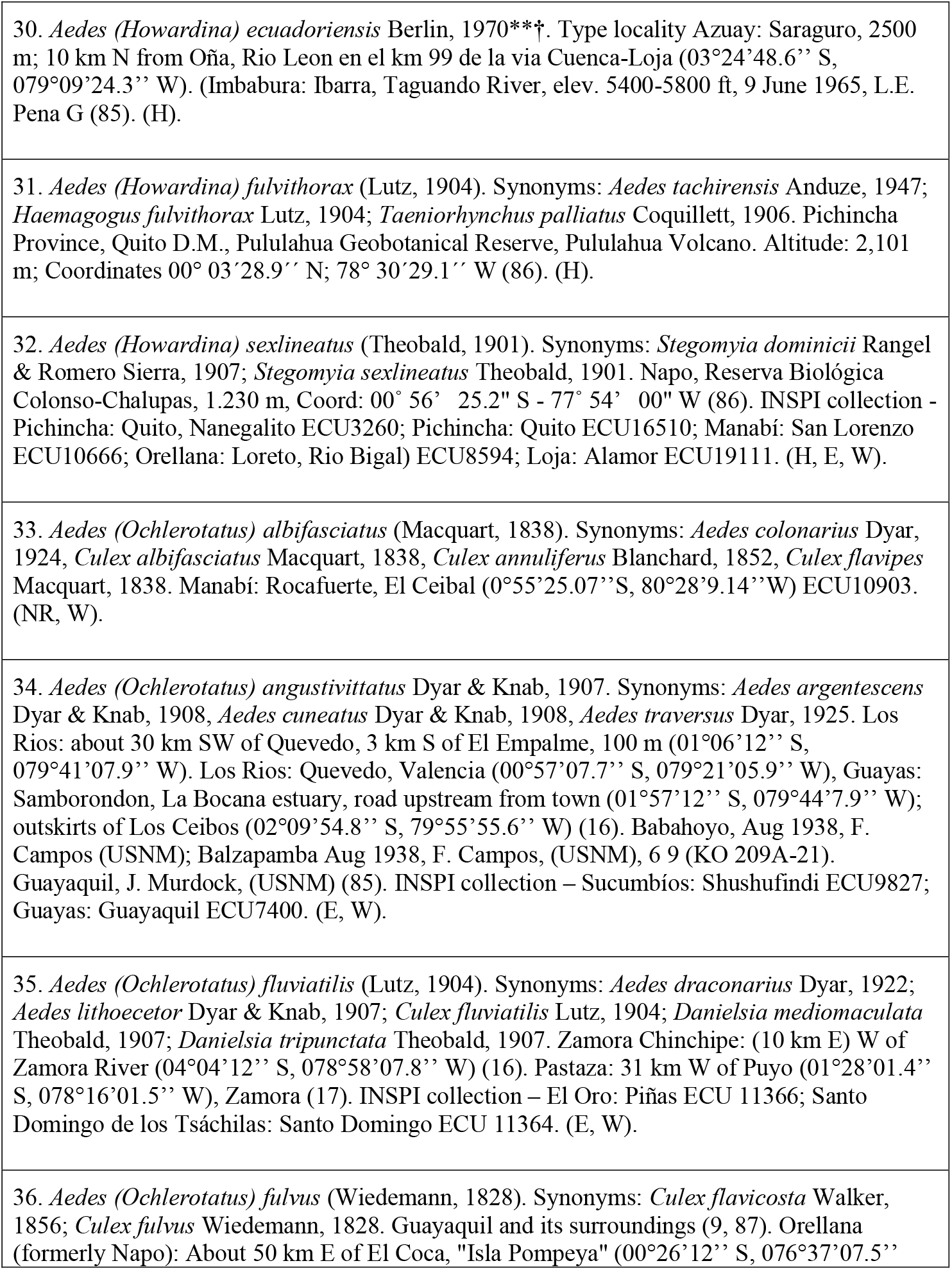
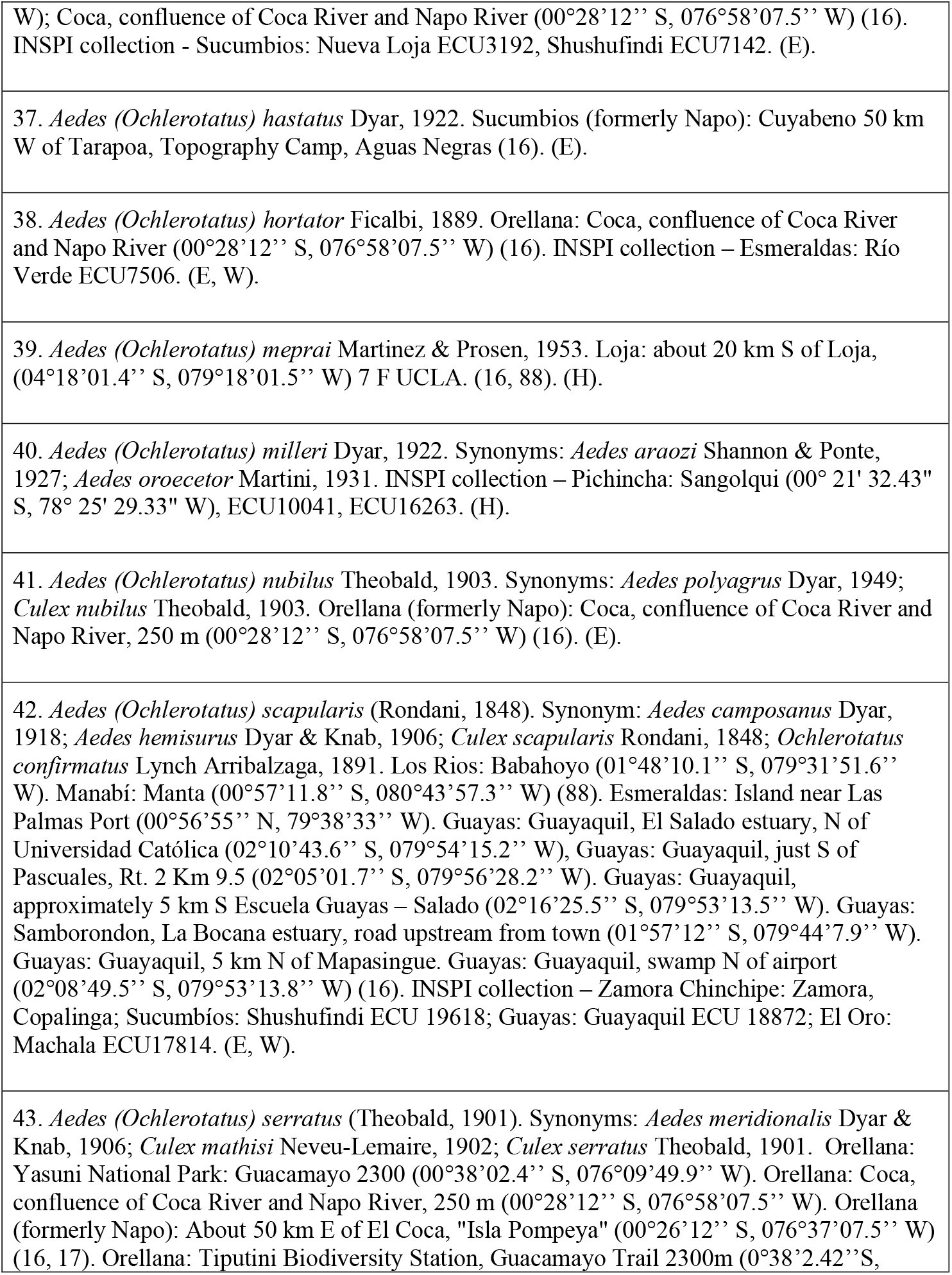
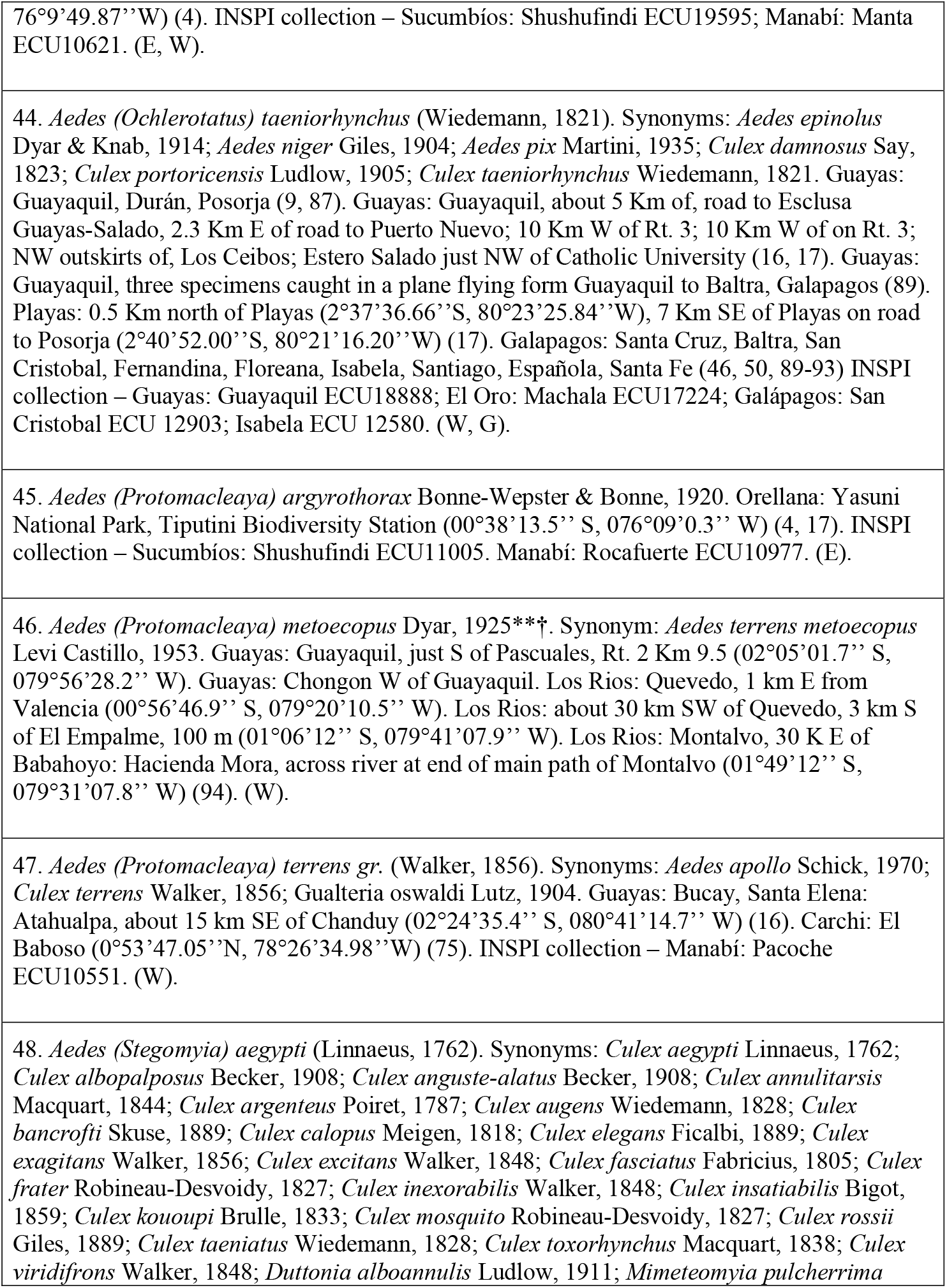
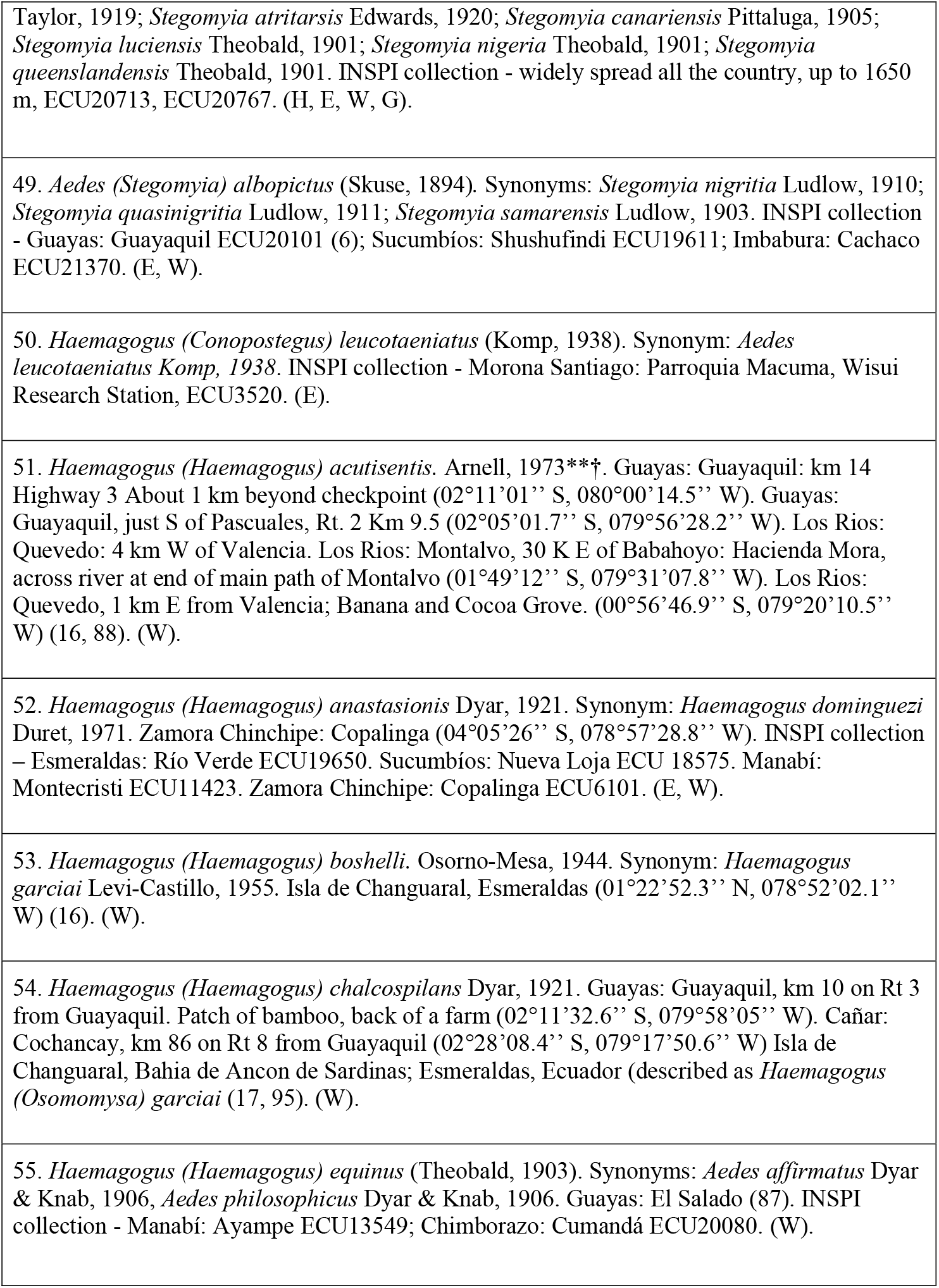
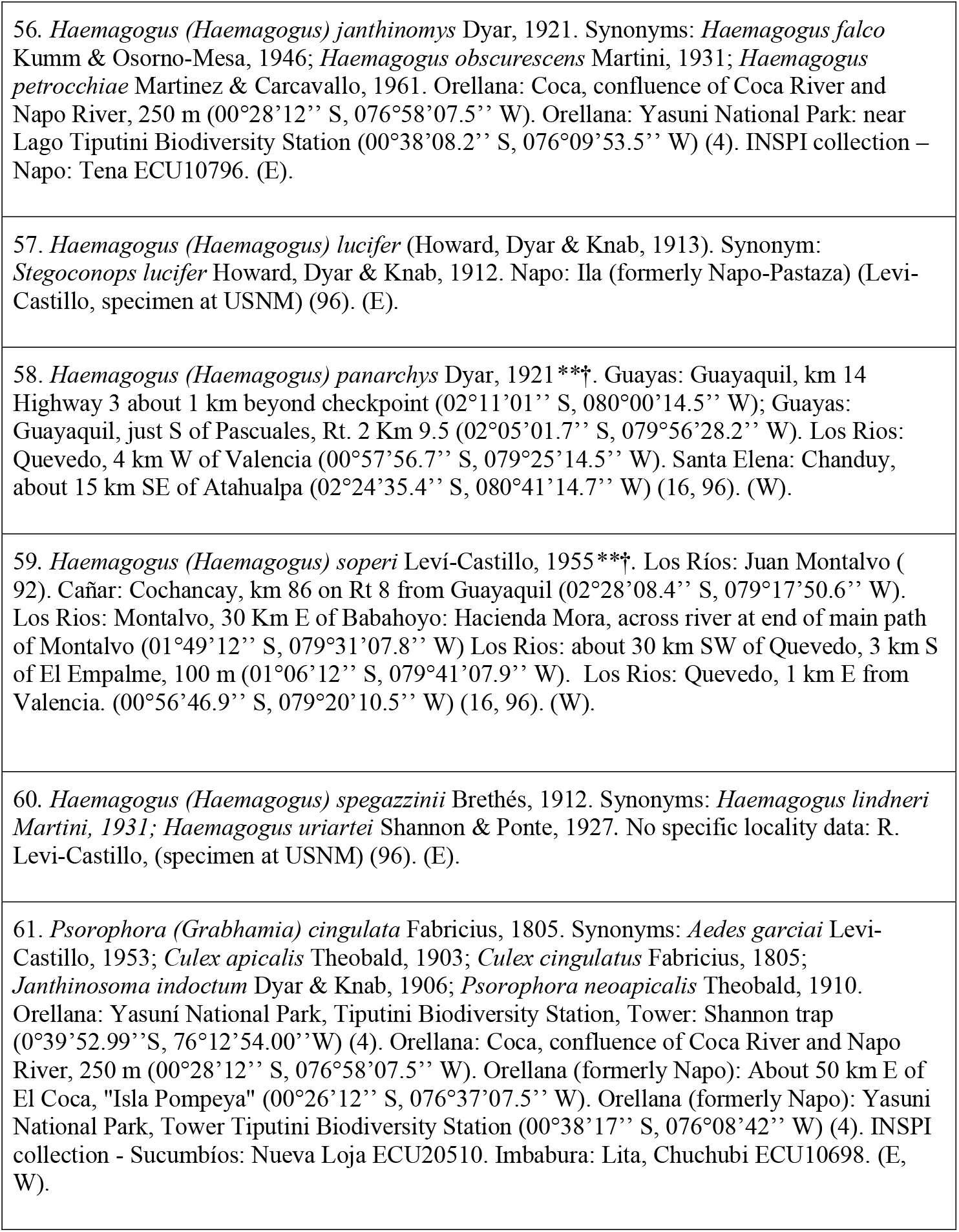
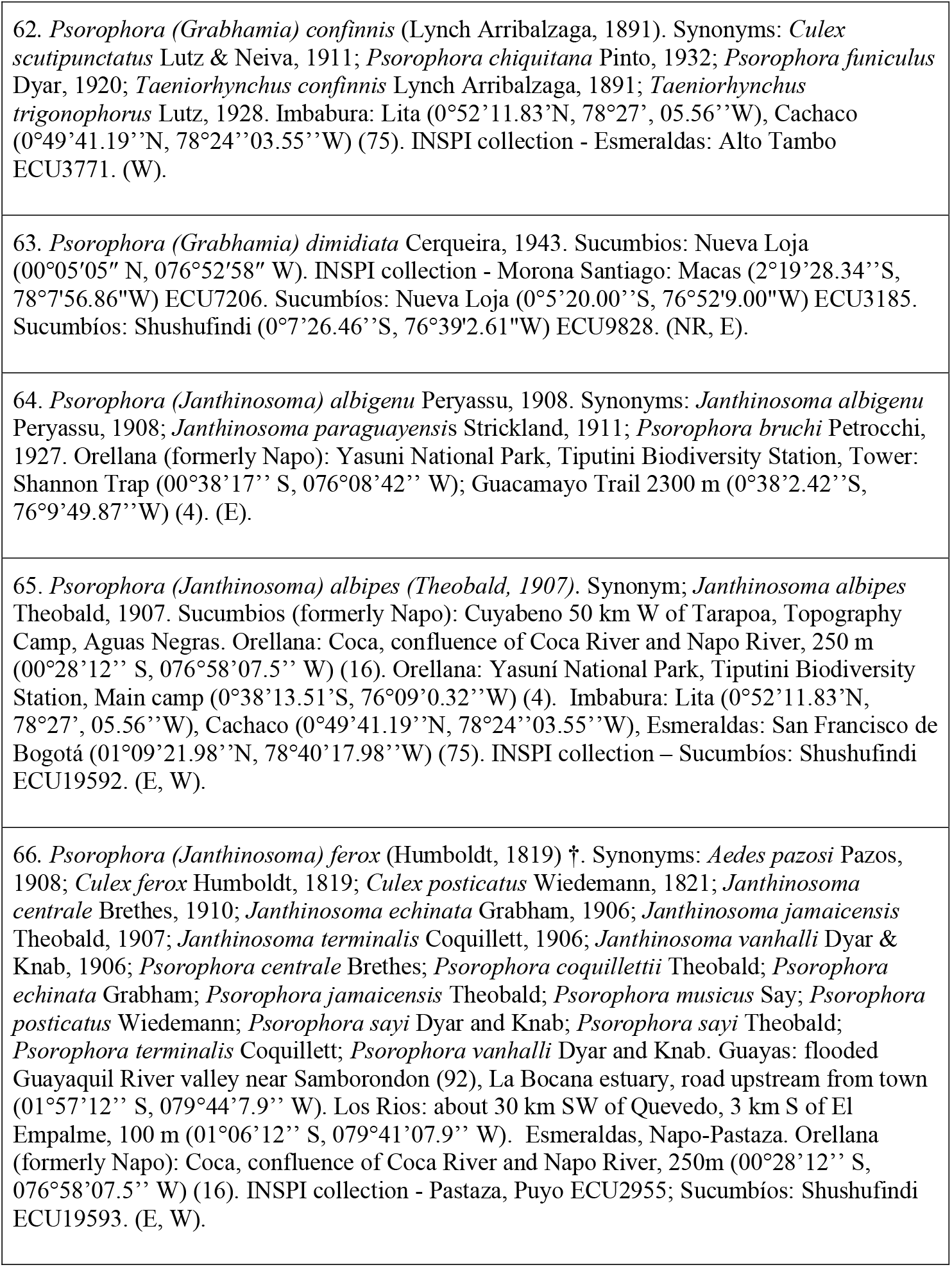
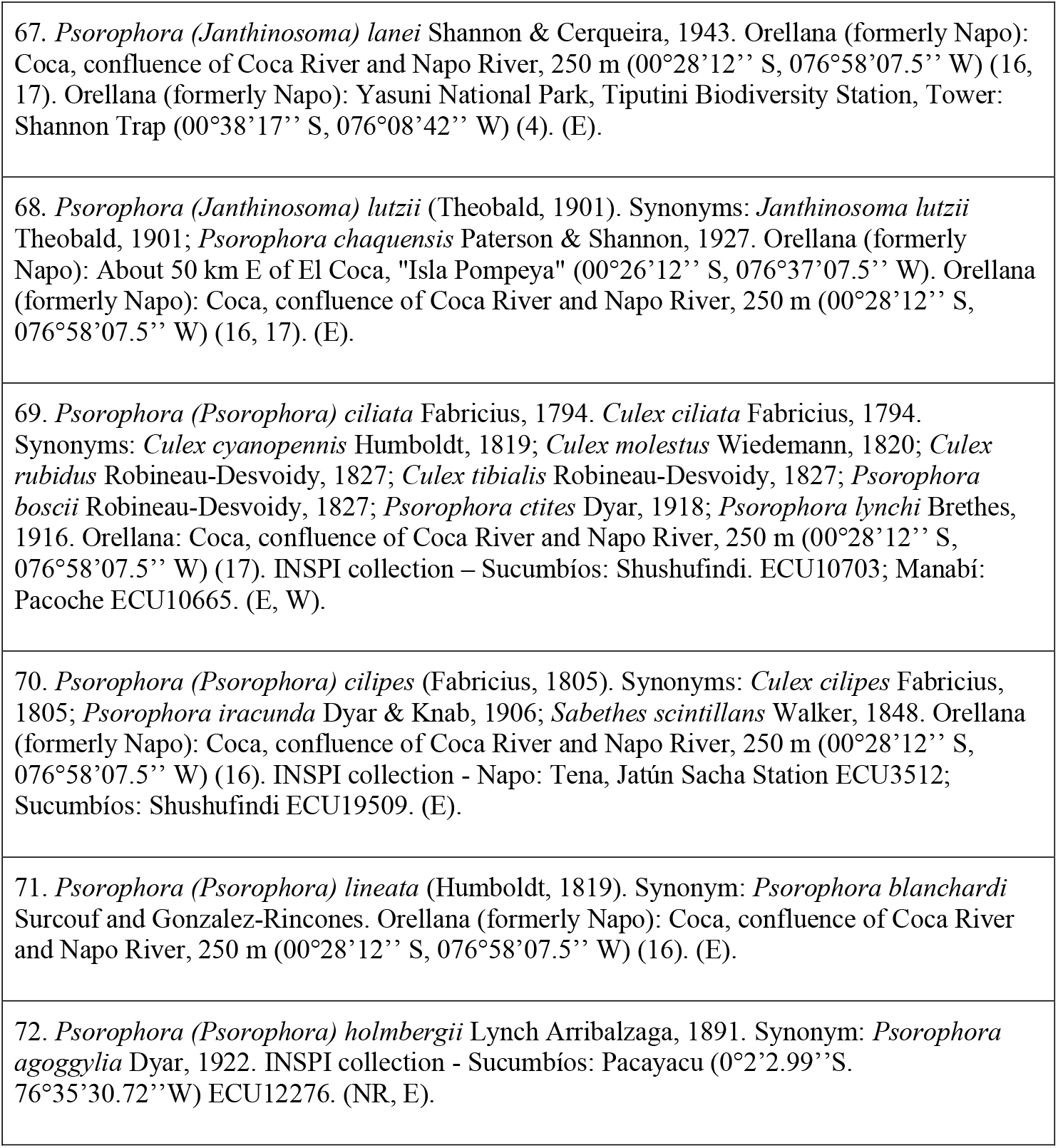
- Subfamily Culicinae. Tribe Aedini; 44 species

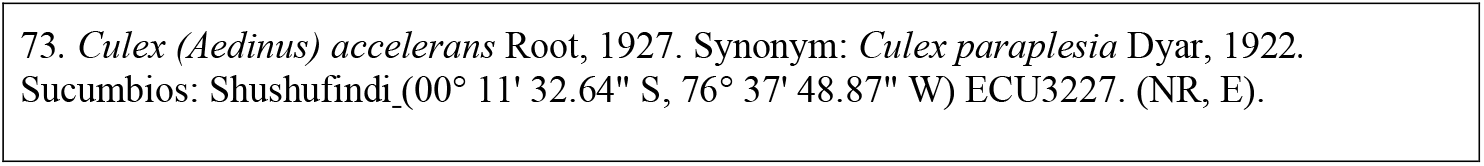
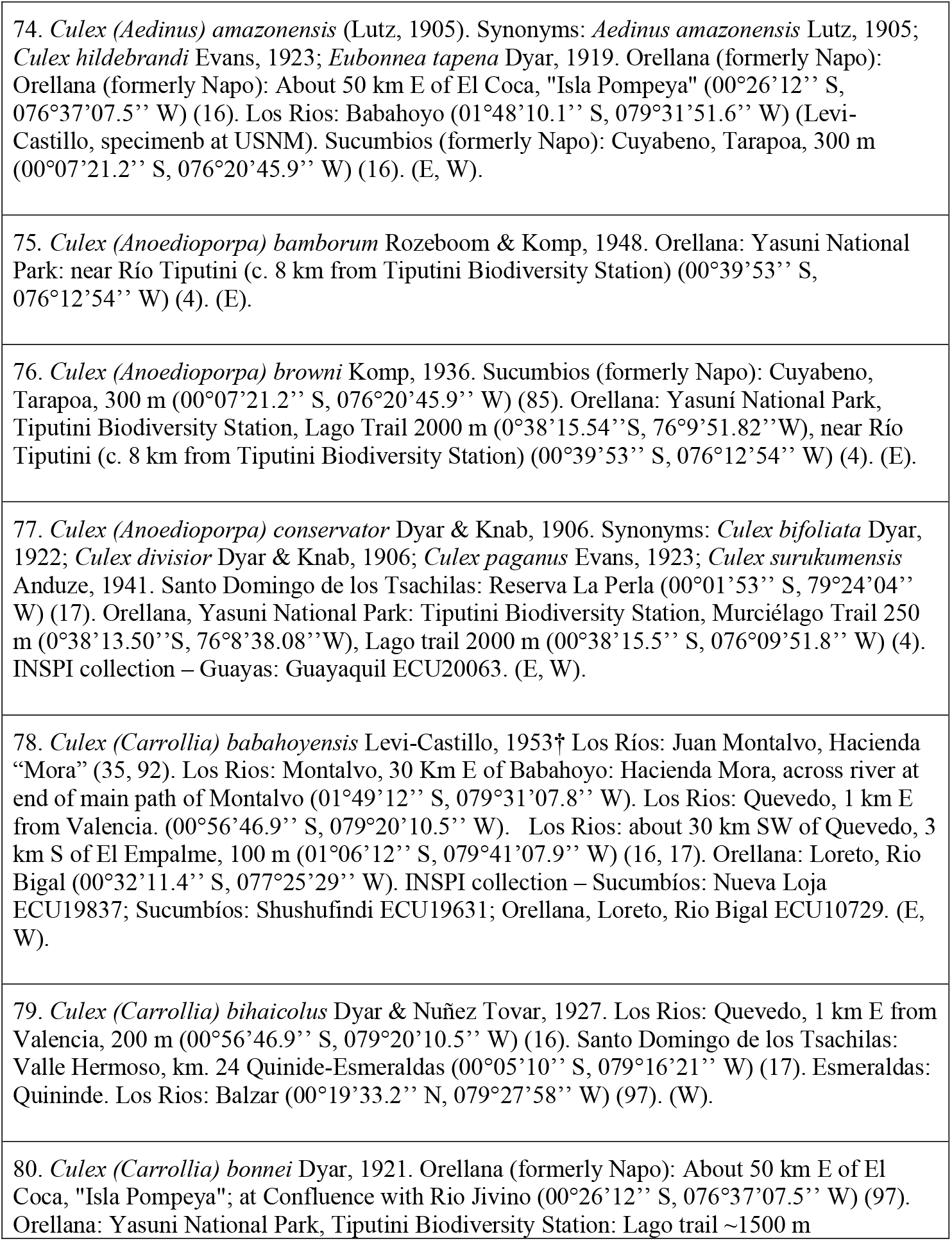
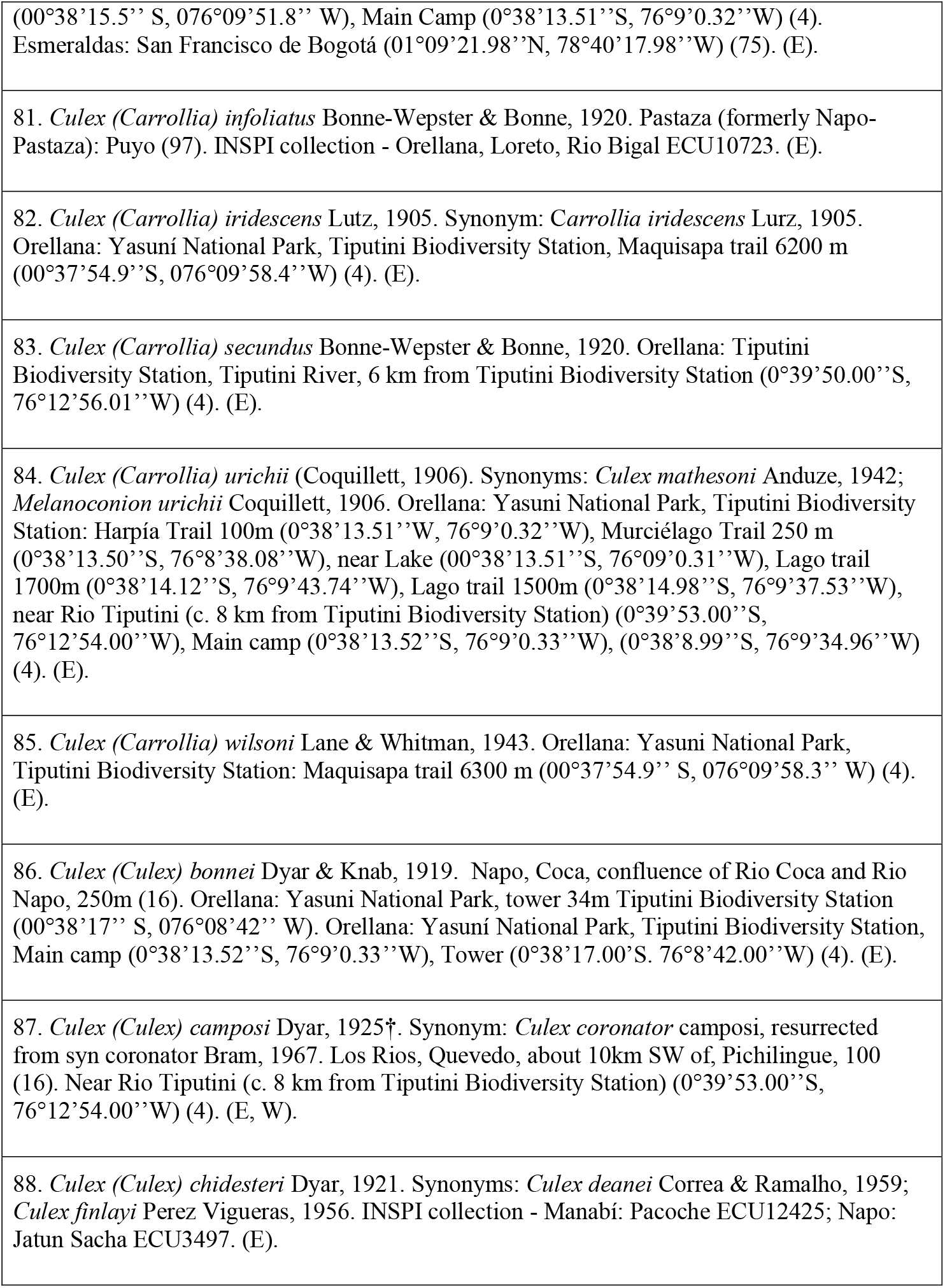
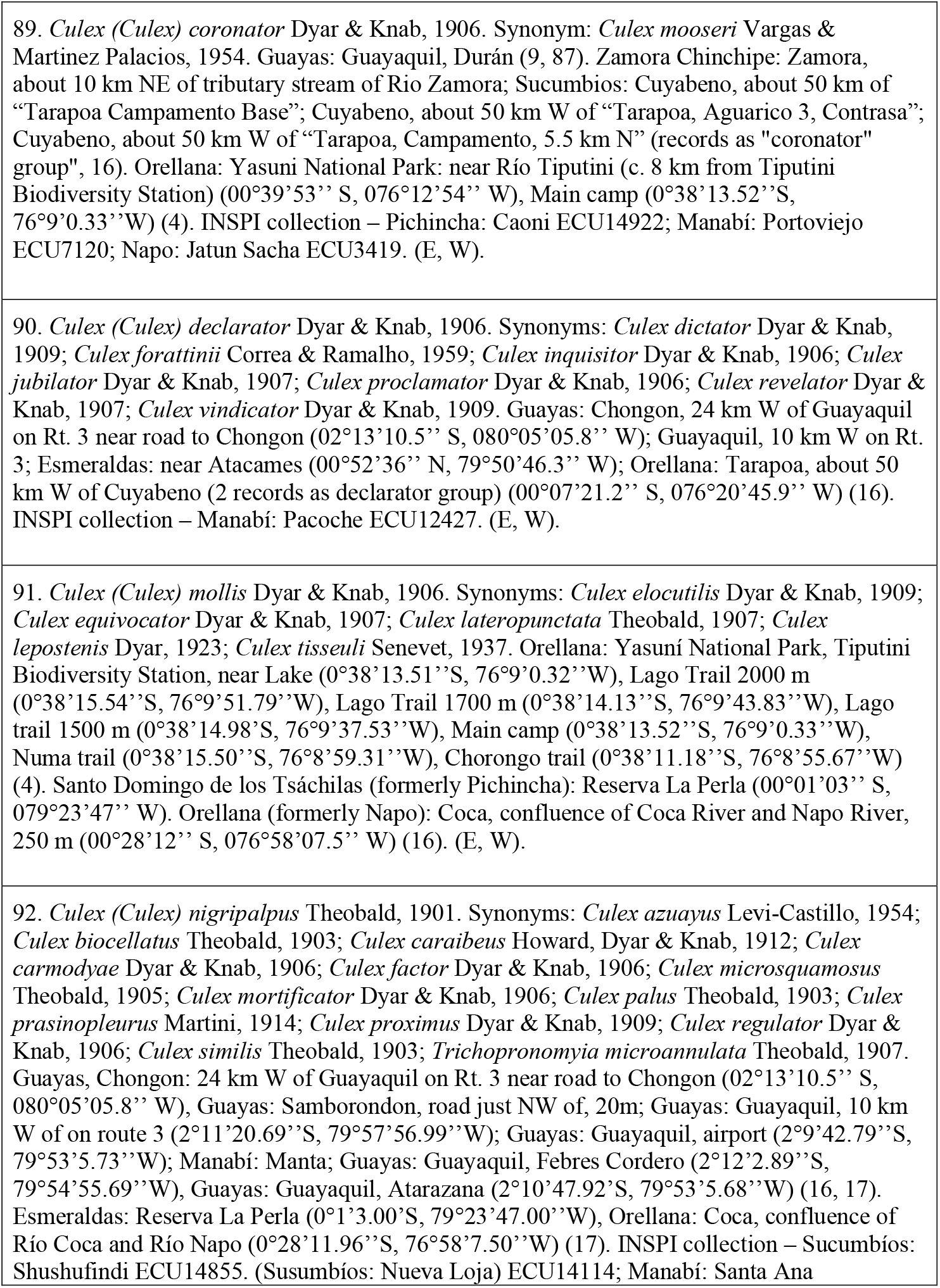
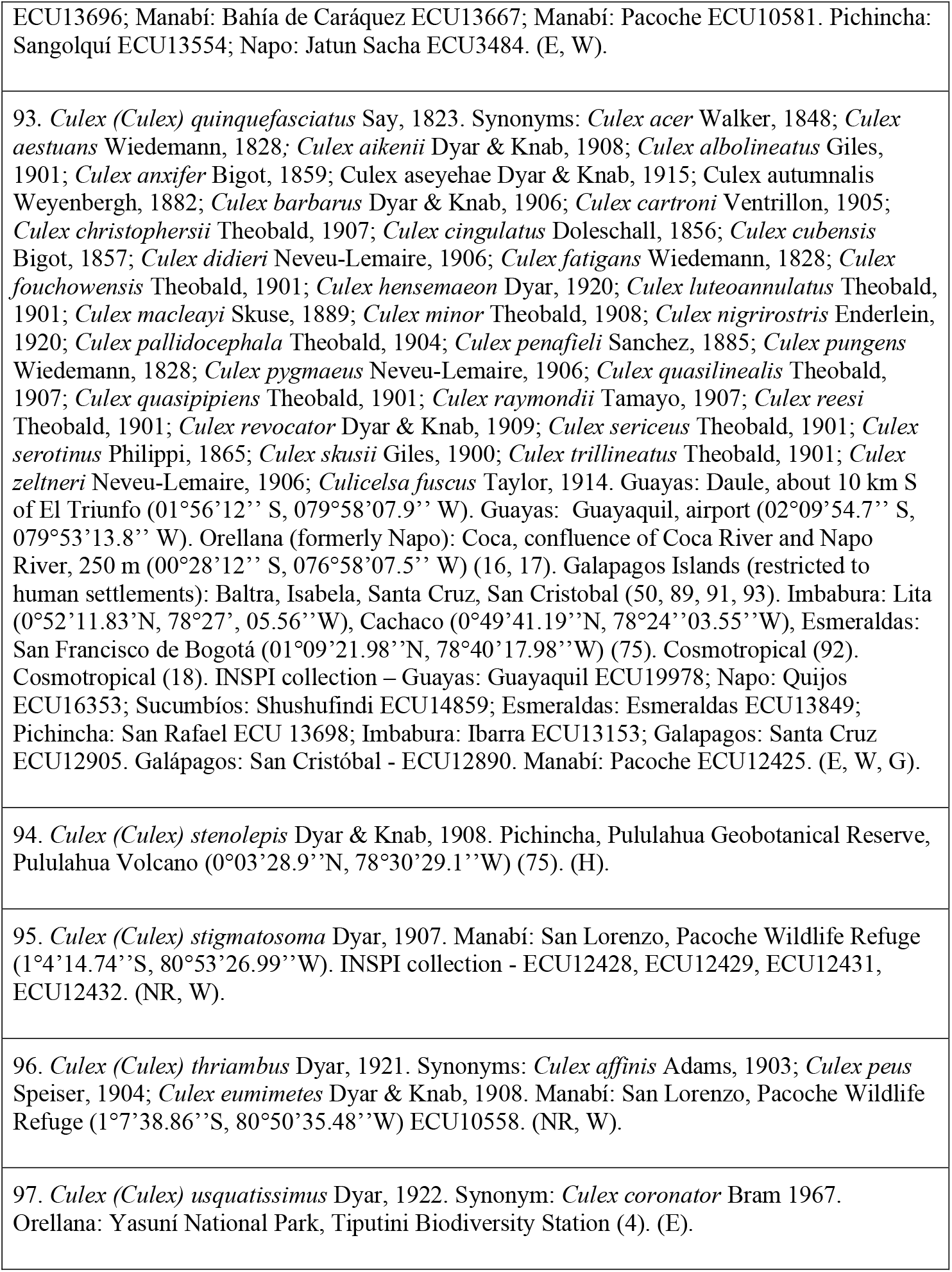
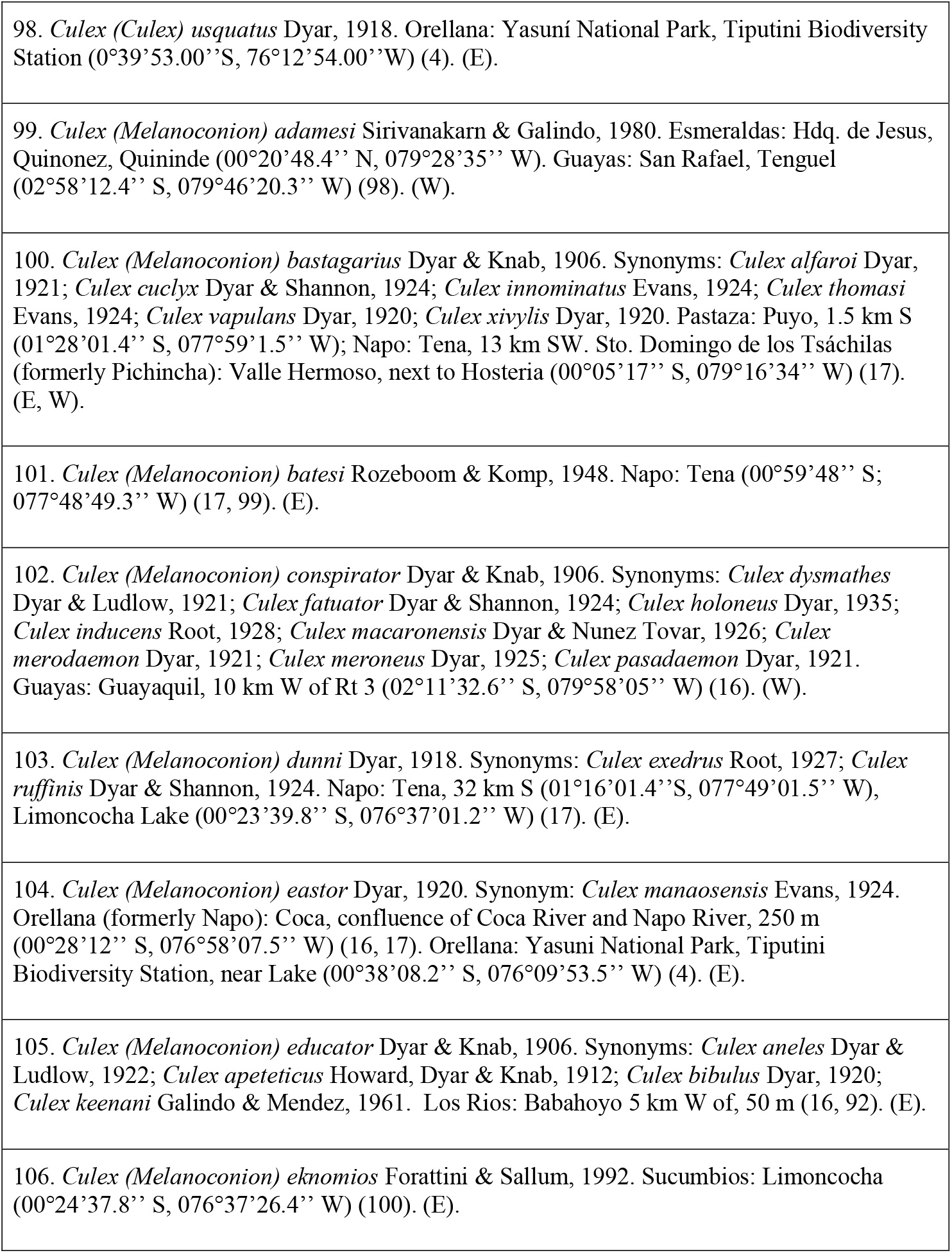
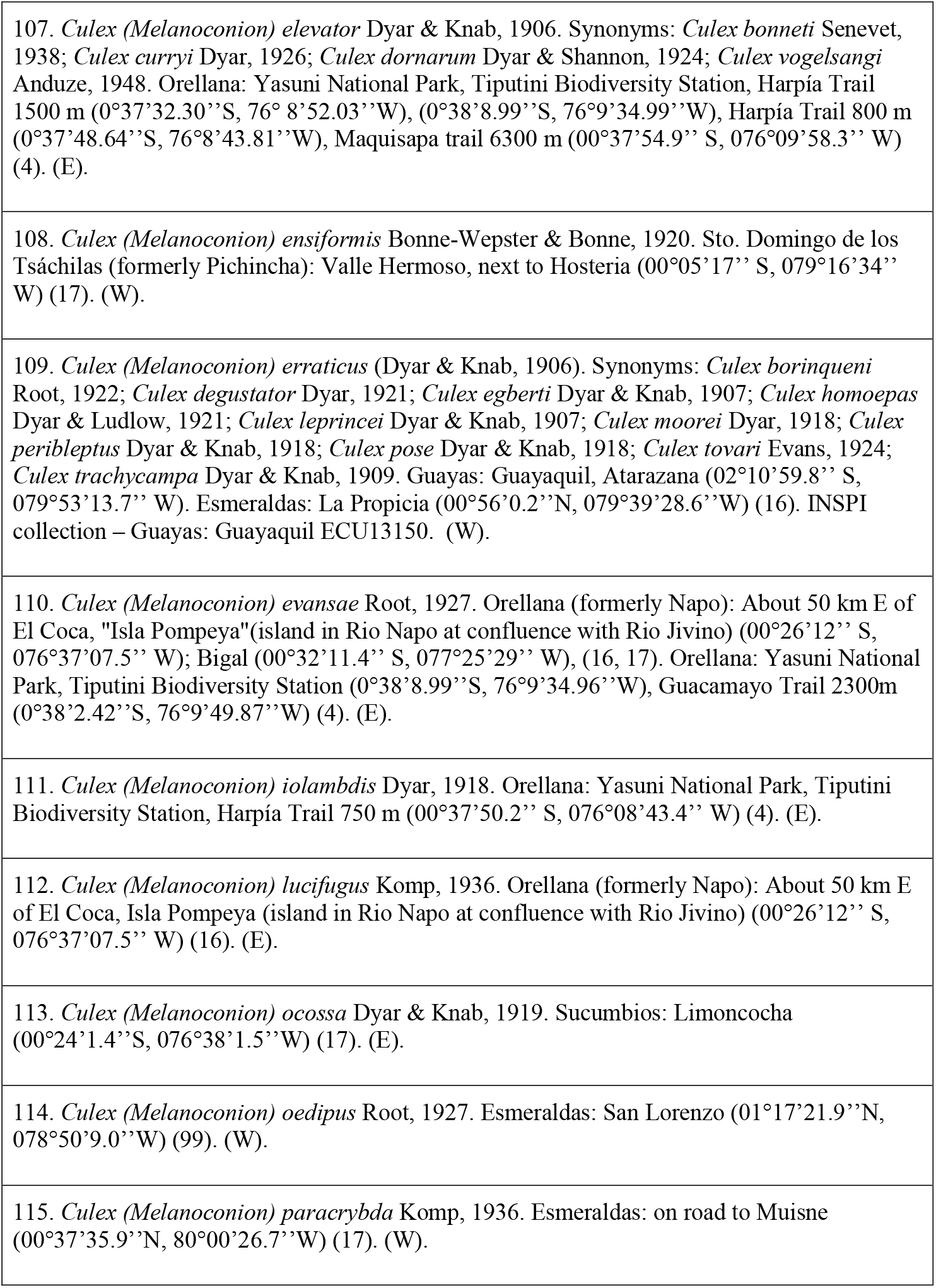
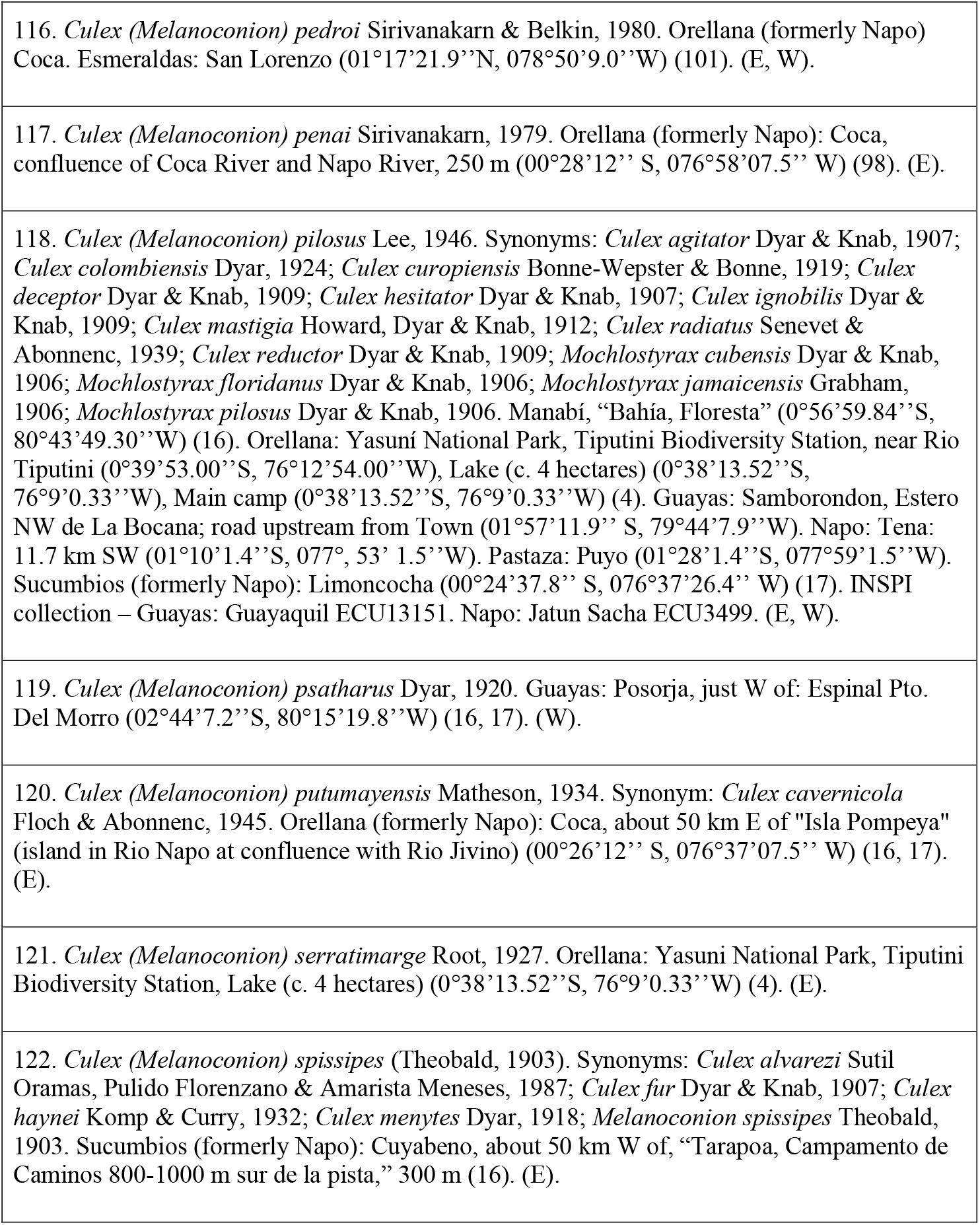
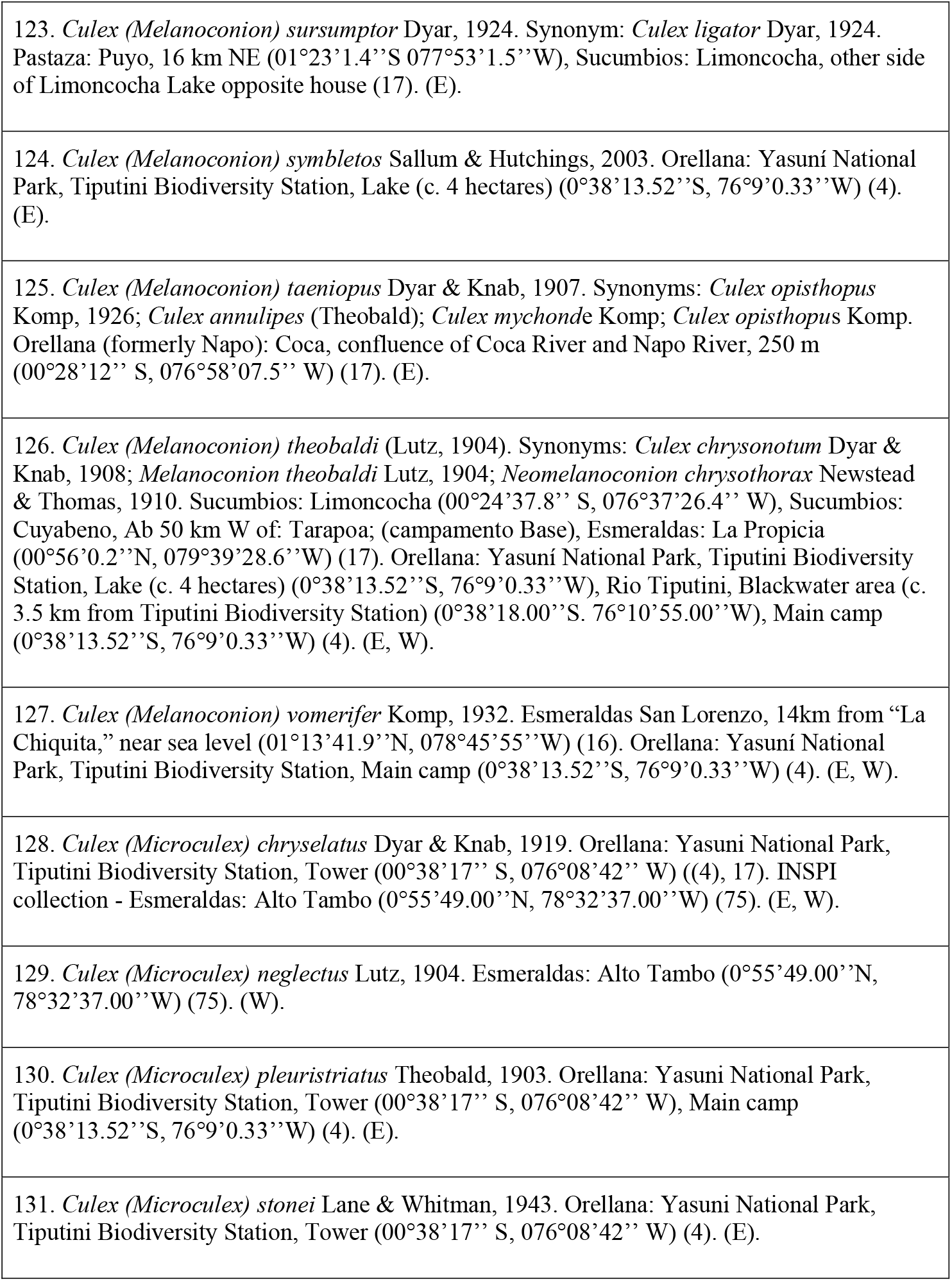
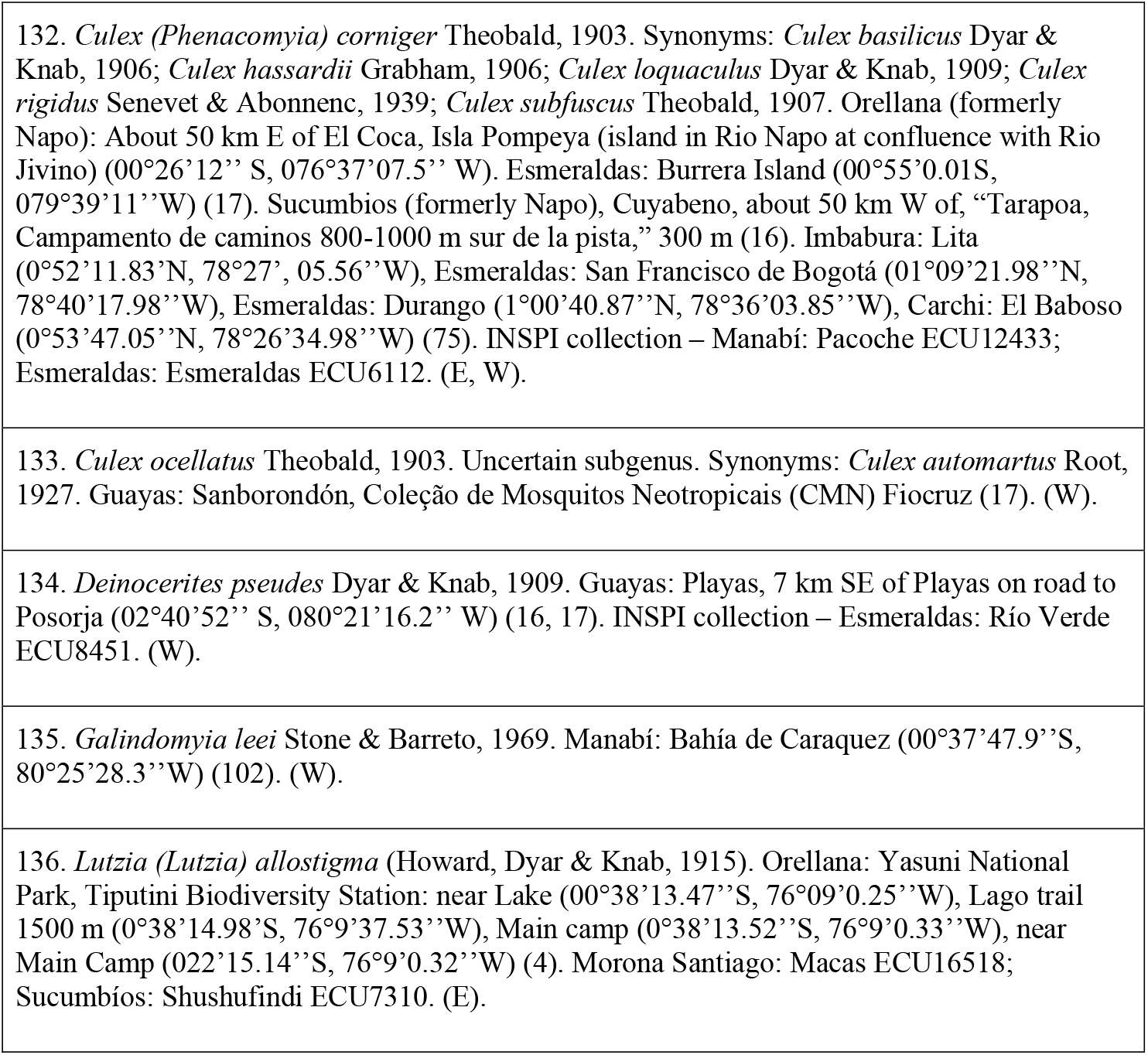
- Tribe Culicini: 64 species

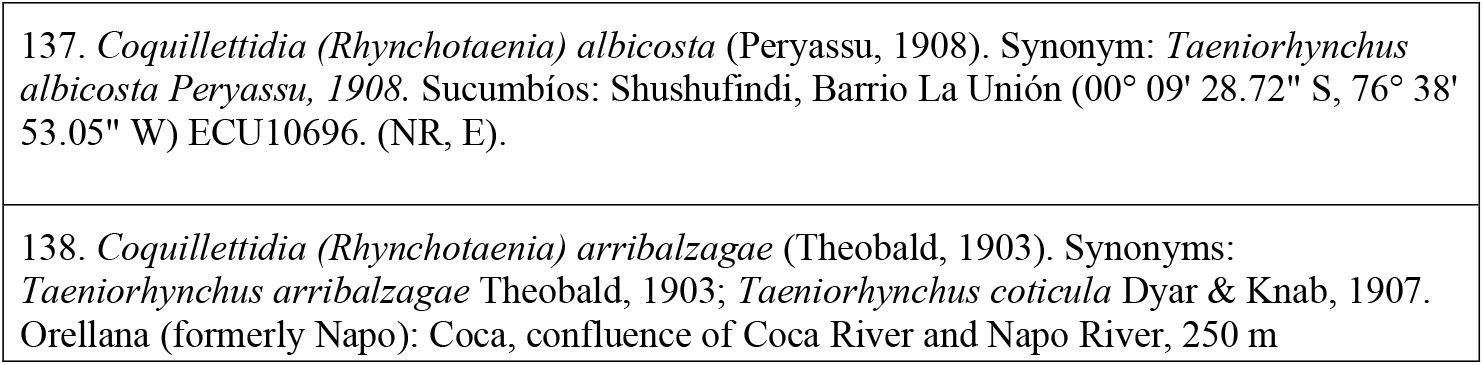
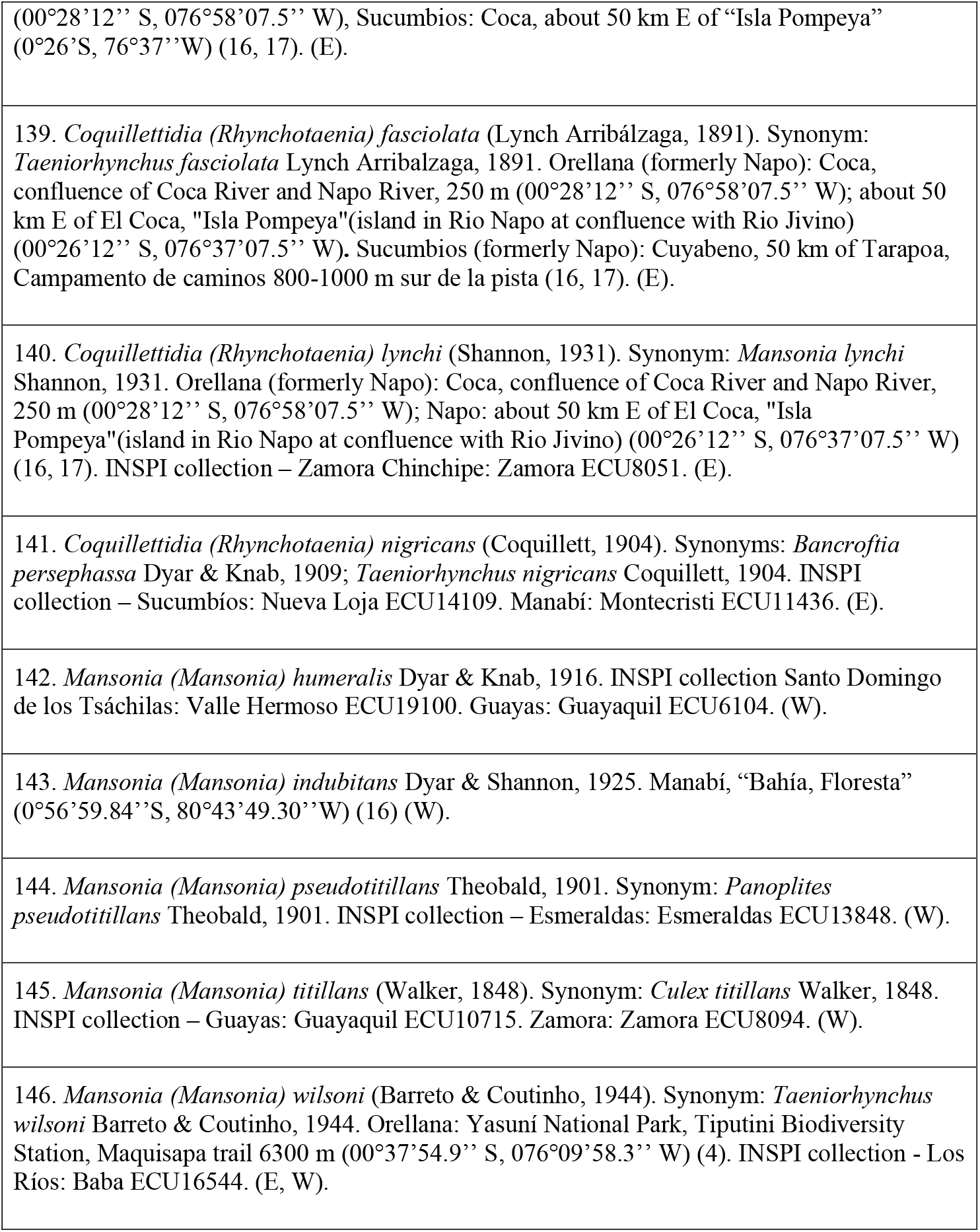
- Tribe Mansoniini:10 species

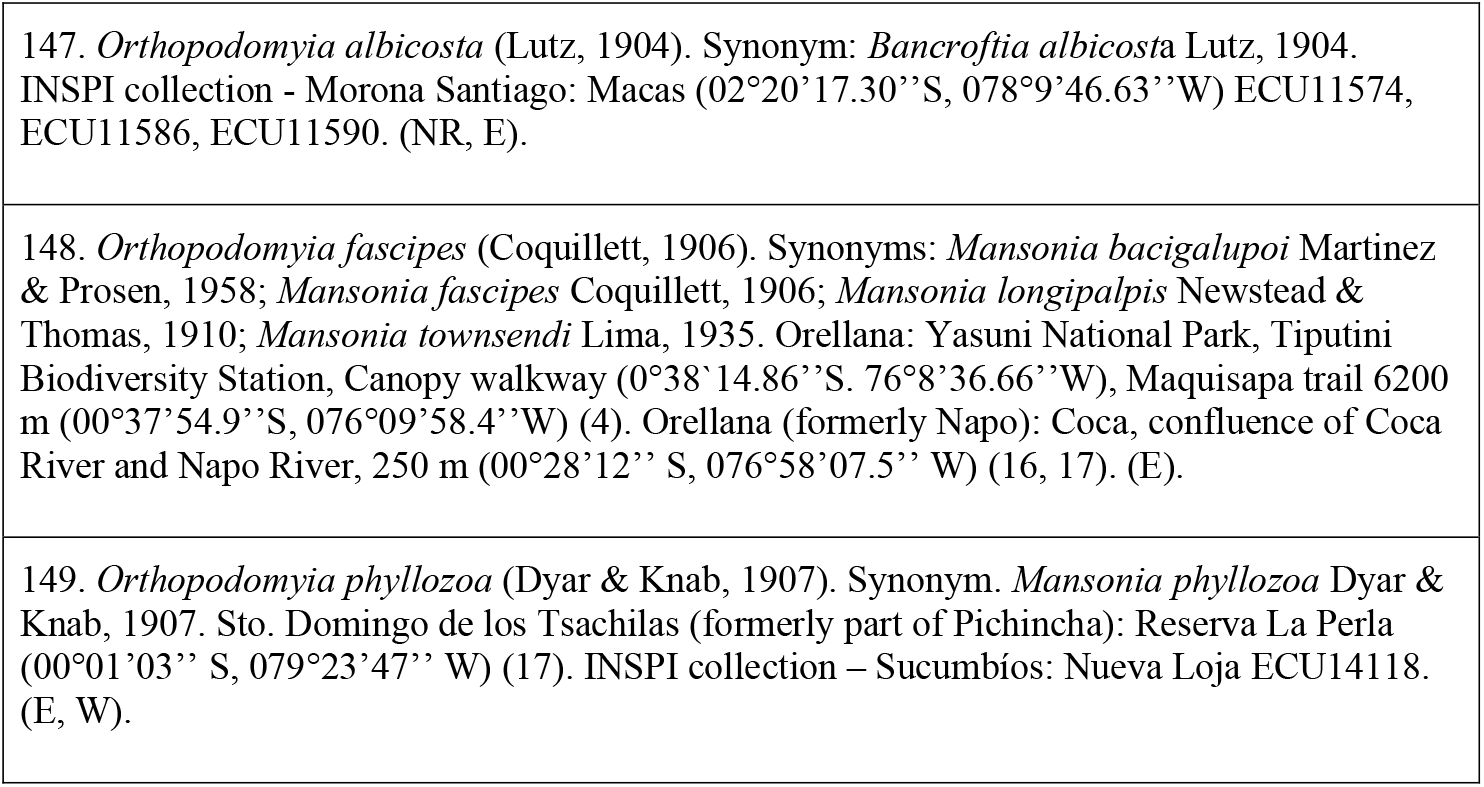
- Tribe Orthopodomyiini: 3 species

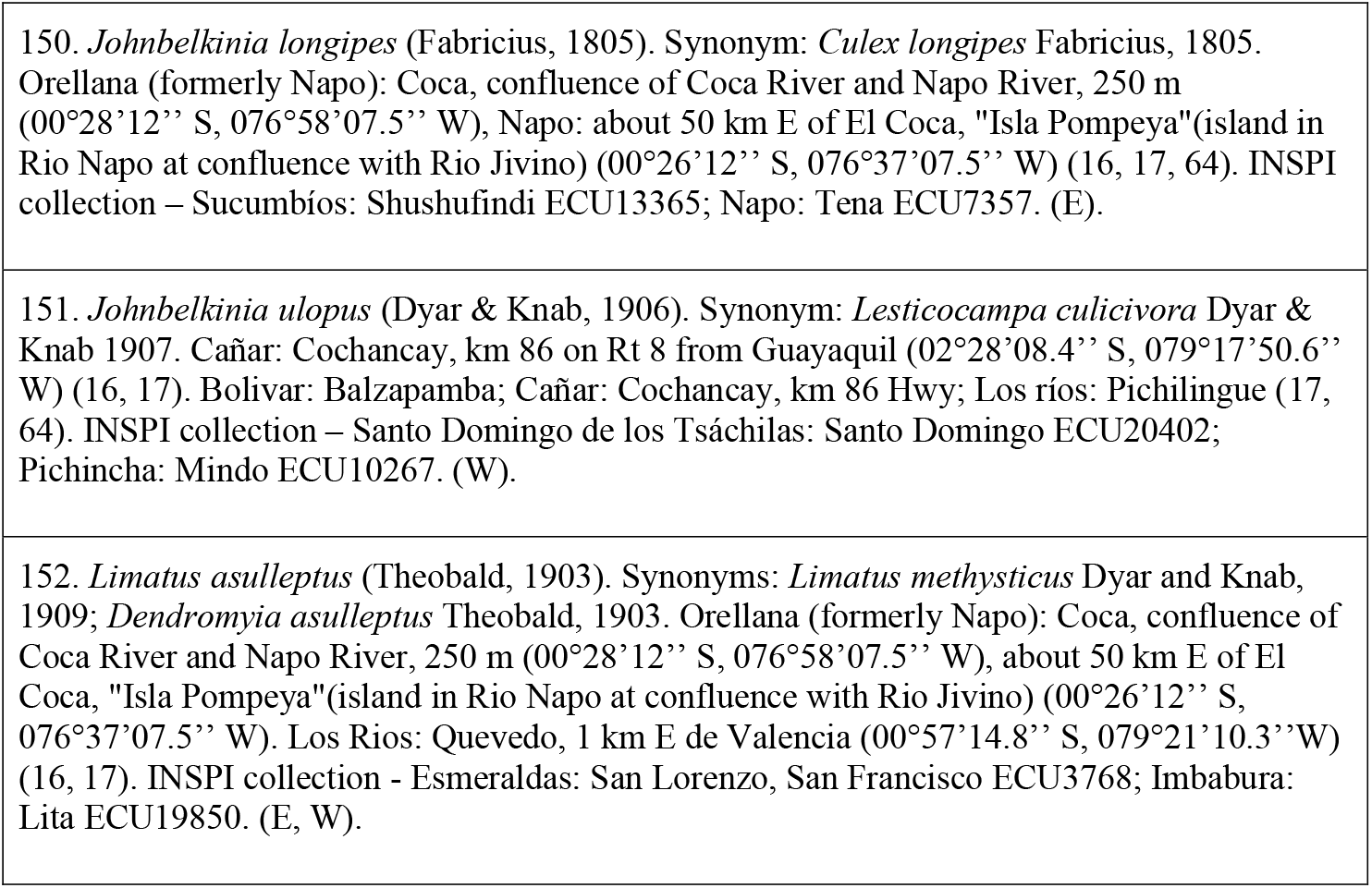
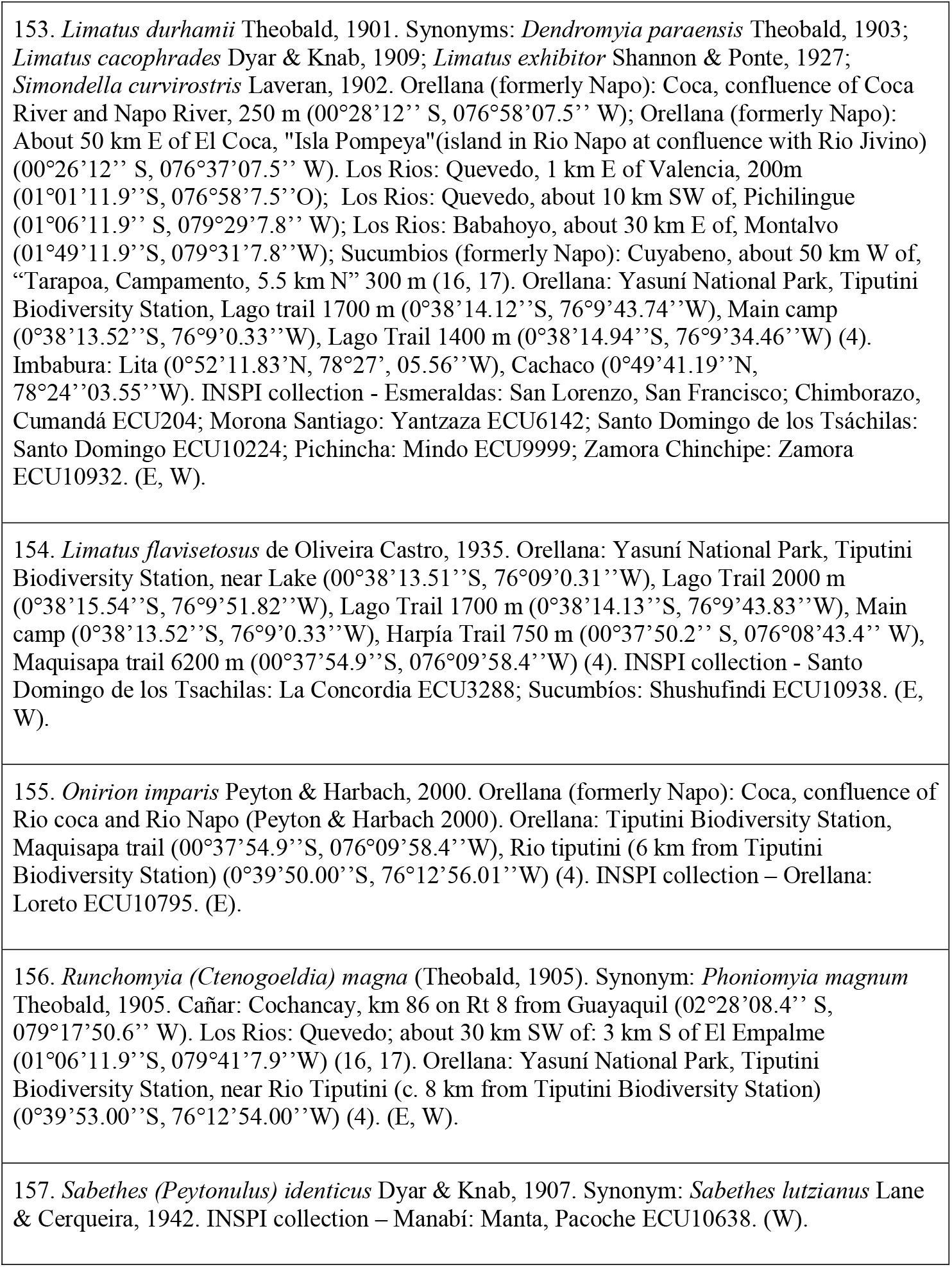
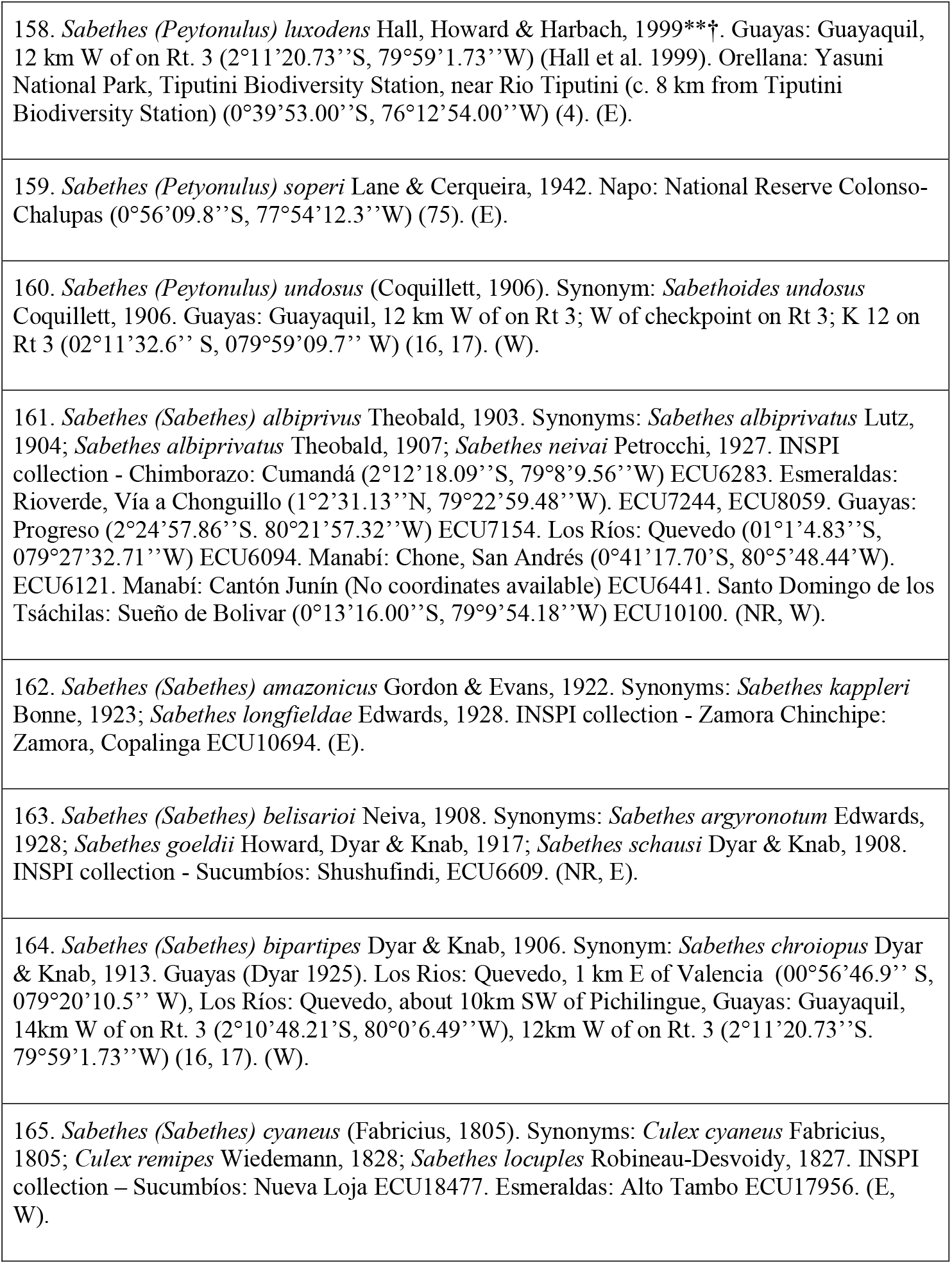
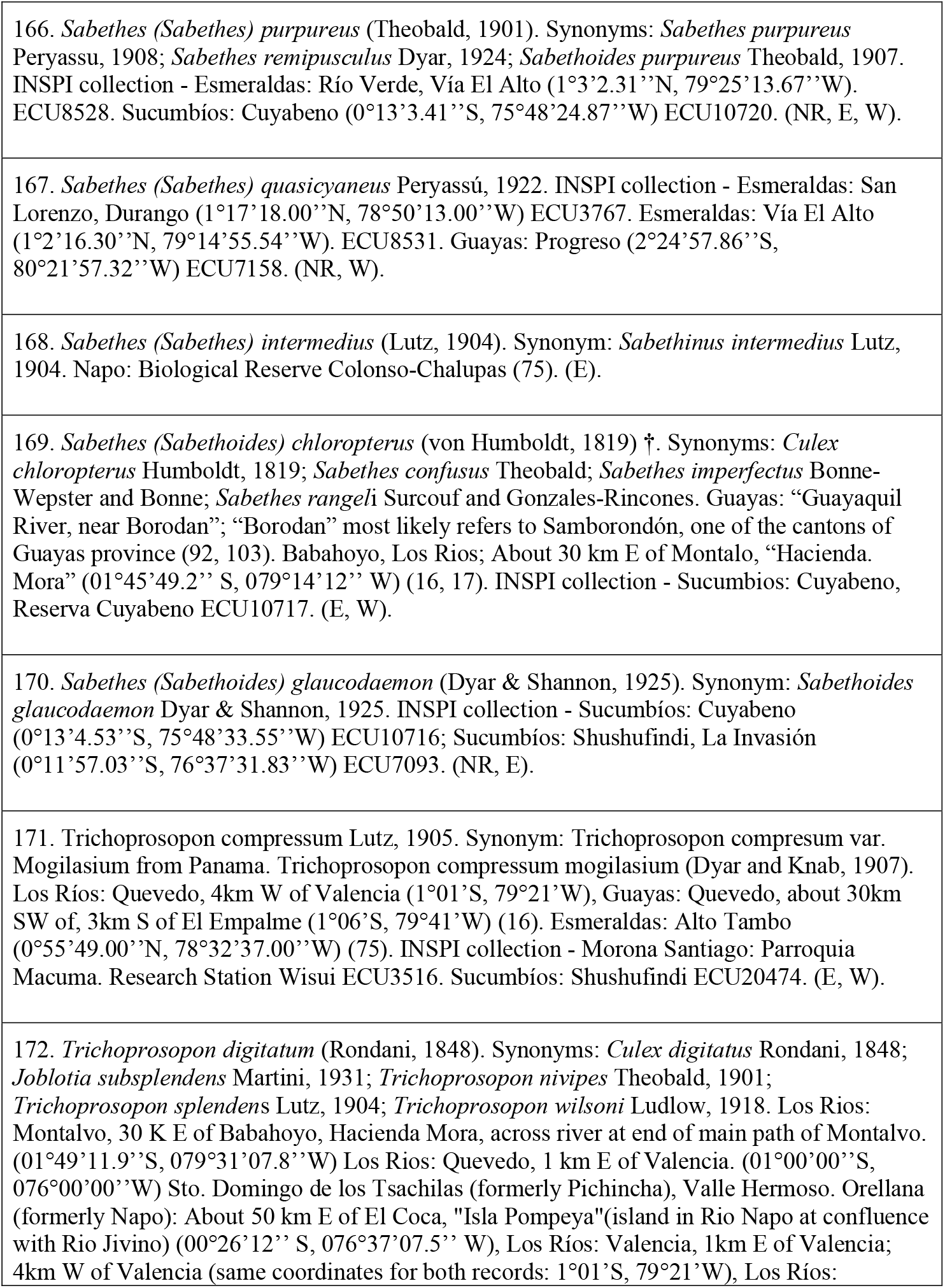
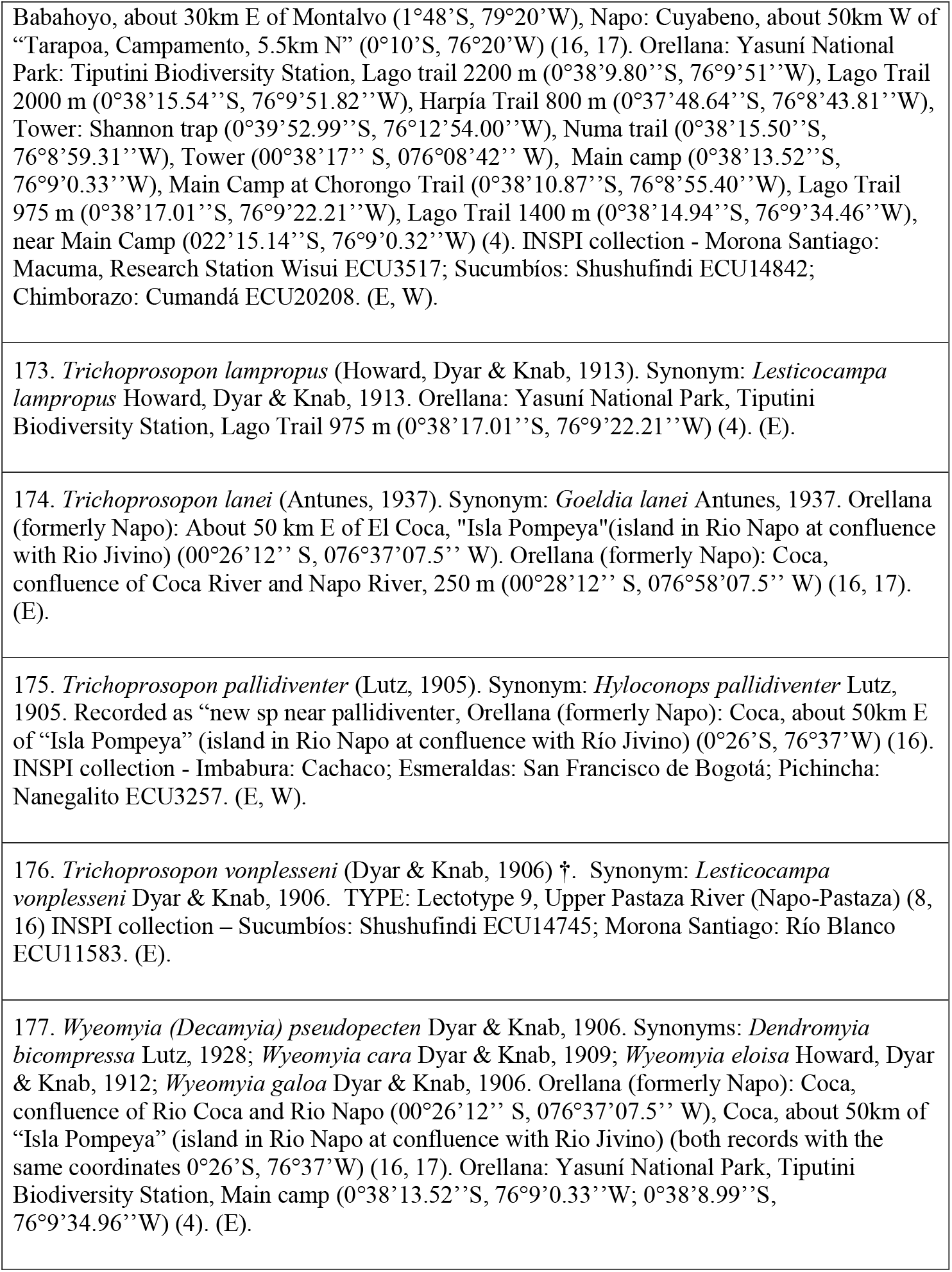
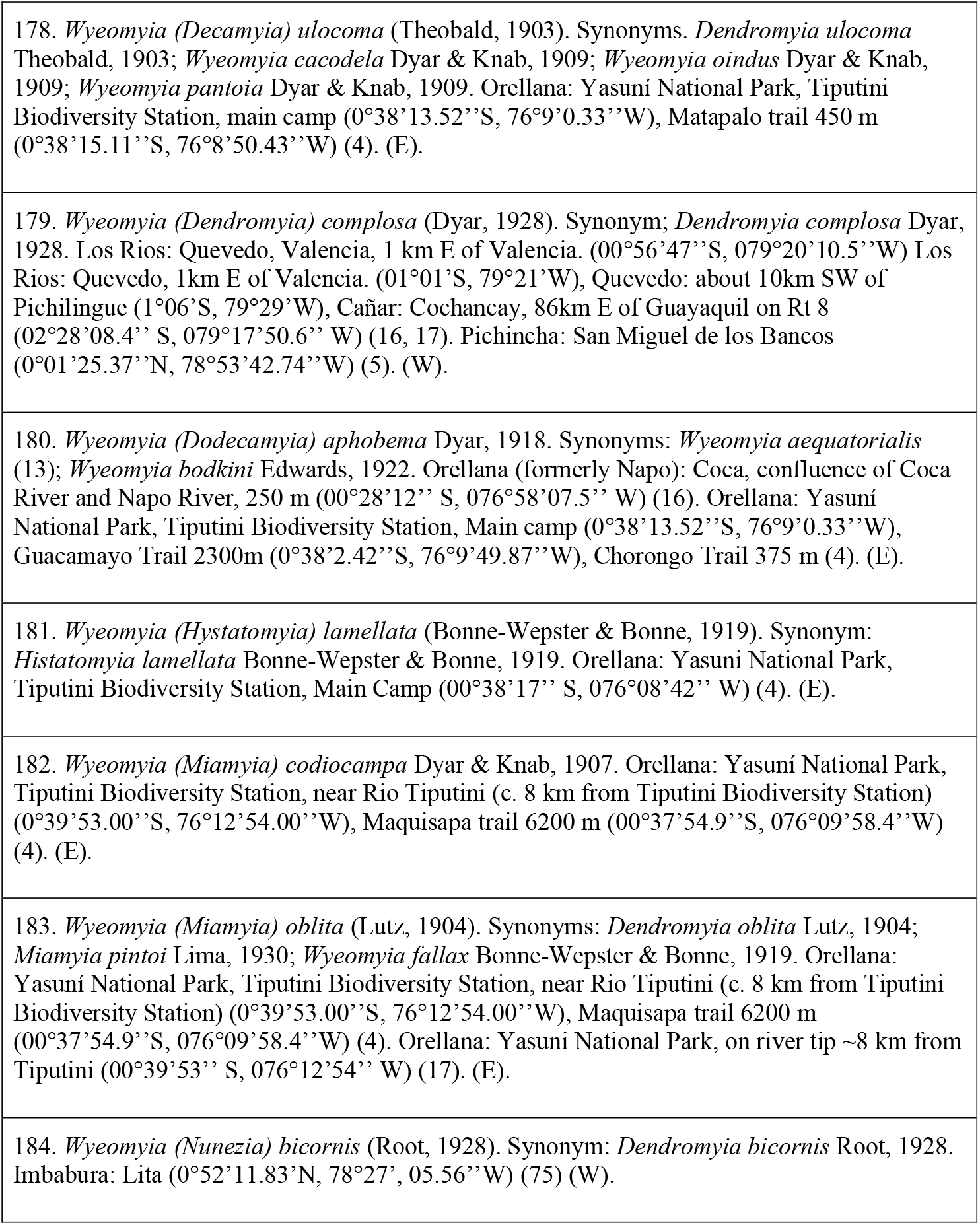
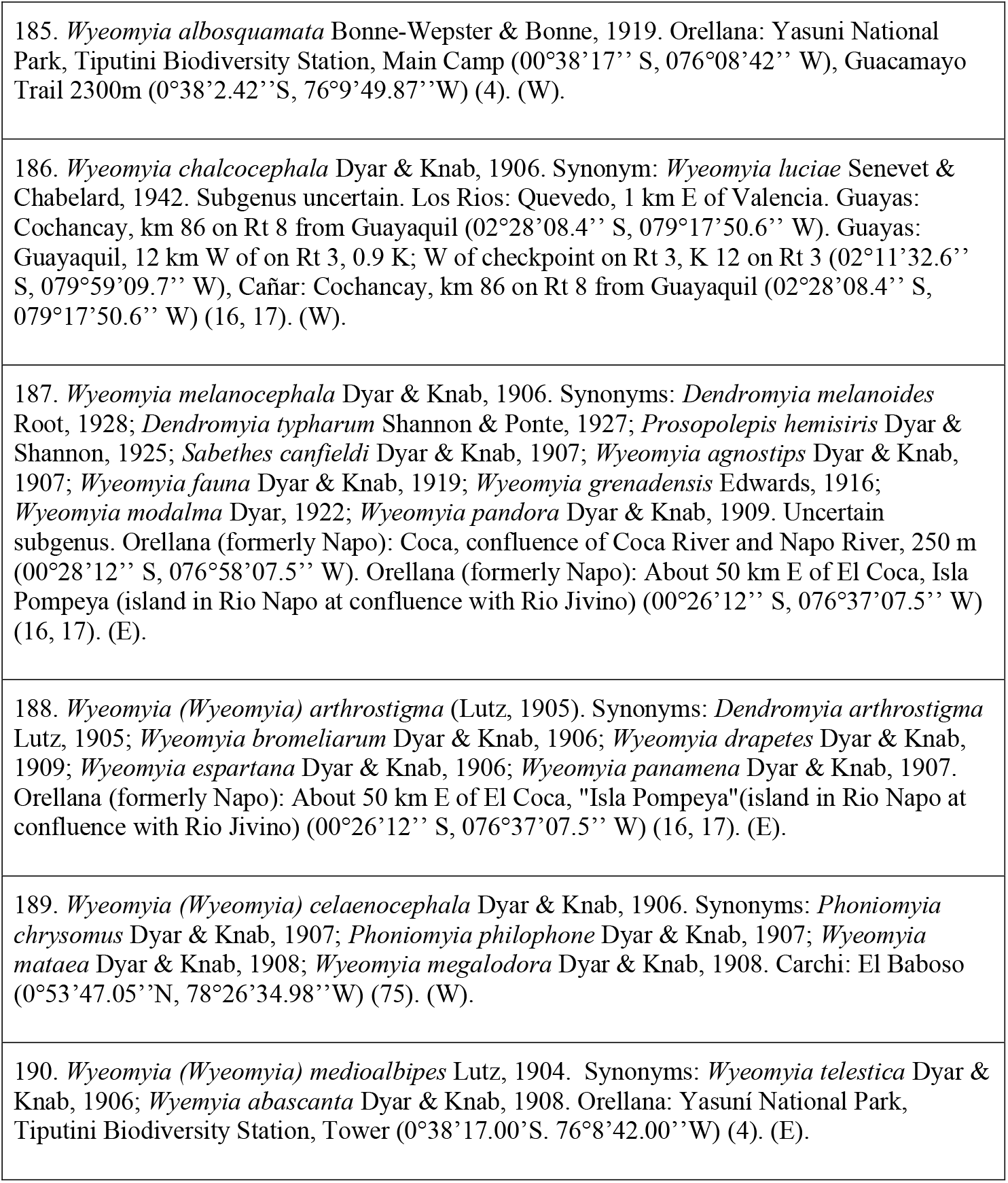
- Tribe Sabethini: 41 species

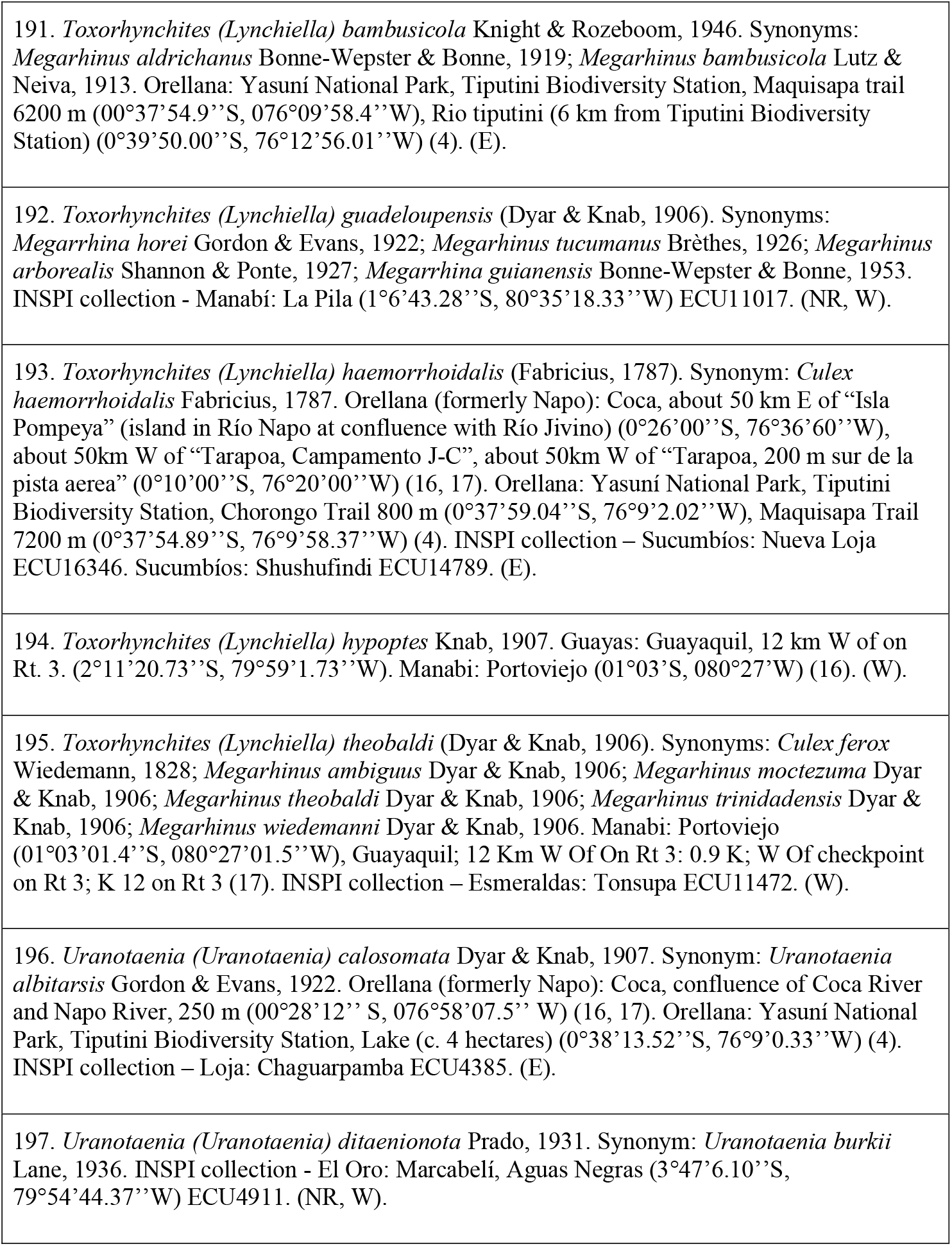
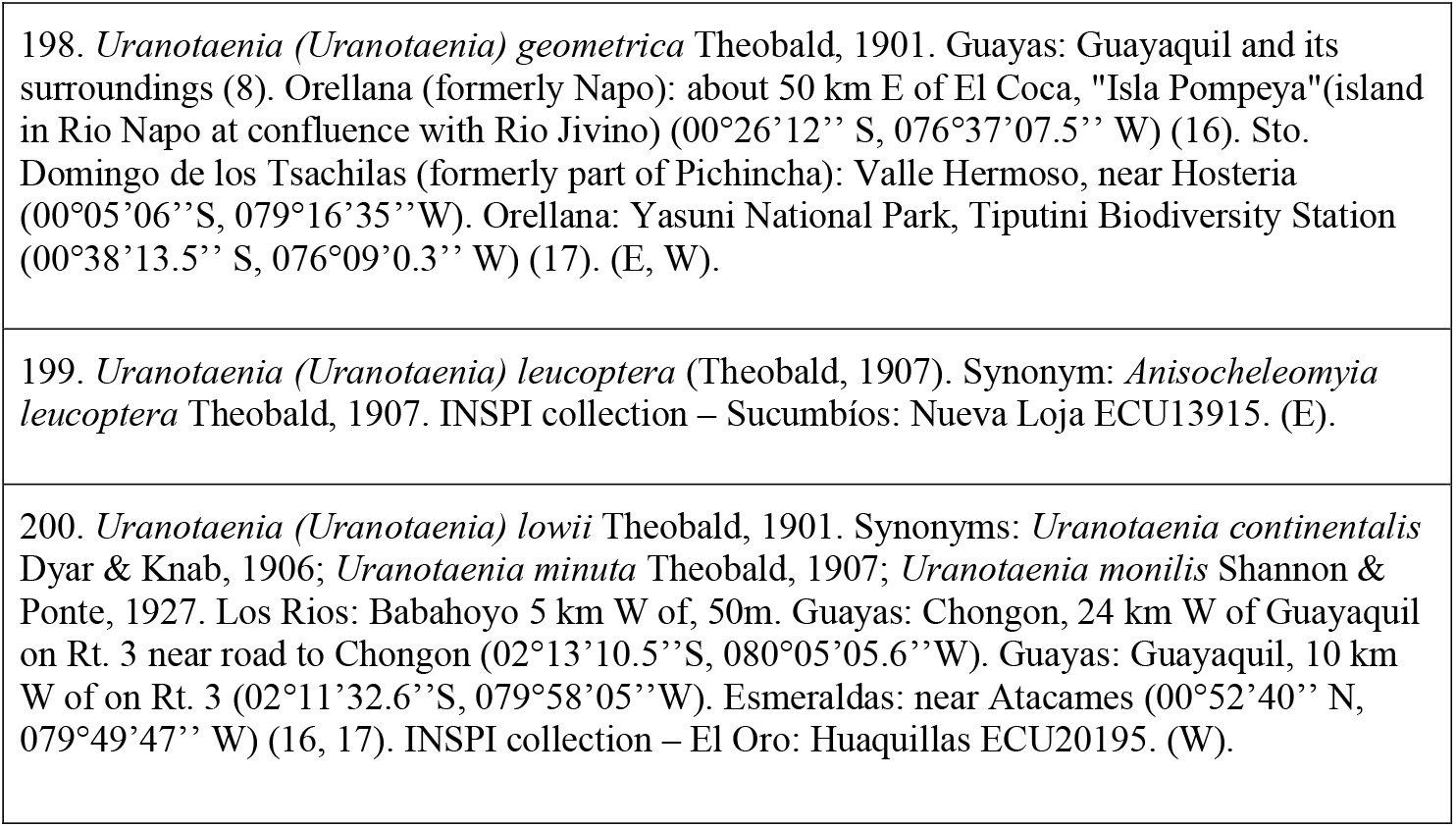
- Tribe Toxorhynchitini: 10 species

## Notes

### Competing Interest Statement

The authors have declared no competing interest.

## References

1. Hooghiemstra H, Wijninga VM, Cleef AM (2006) The paleobotanical record of Colombia: implications for biogeography and biodiversity. Ann. Missouri Bot. Gard. 93: 297–325.

2. Holdridge LRW, Grenke W, Hatheway H, Liang H, and Tosi JA (1971) Forest environments in tropical life zones: a pilot study. Pergamon Press. Oxford, UK. 747.

3. Cañadas L (1983) El mapa bioclimático del Ecuador. Banco Central del Ecuador. Quito.

4. Linton YM, Pecor JE, Porter CH, Mitchell LB, Garzón-Moreno A, Foley DH, Pecor DB, Wilkerson RC (2013) Mosquitoes of eastern Amazonian Ecuador: biodiversity, bionomics and barcodes. Mem Inst Oswaldo Cruz. 108: 100–109.

5. Navarro JC, Ponce P, Cevallos V (2013) Dos nuevos registros de vectores potenciales de Fiebre Amarilla selvática y Mayaro para el Ecuador. Bol Malariol y Salud Ambient. 53: 77–81.

6. Ponce P, Morales D, Argoti A, Cevallos VE (2018) First Report of *Aedes (Stegomyia) albopictus* (Skuse) (Diptera: Culicidae), the Asian Tiger Mosquito, in Ecuador. J Med Entomol. 55(1): 248–249.

7. Harbach RE (2013) Mosquito Taxonomic Inventory. Available at: http://mosquito-taxonomic-inventory.info/.

8. Belkin JN, Schick RX, Galiudo P, Aitkeu THG (1965) Mosquito studies (Diptera: Culicidae) 1. A project for a systematic study of the mosquitoes of Middle America. II. Methods for collection, rearing and preservation of mosquitoes. Co&b. Am Entomol Inst. 1(2):1–78.

9. Campos FR (1925) Estudios Biológicos sobre los mosquitos de Guayaquil y alrededores. Rev Col Nac Vicente Rocafuerte. 7: 21–22.

10. Campos FR (1928) Algo sobre cromotaxia de las larvas del mosquito *malárico Anopheles tarsimaculatus* Goeldi. Rev Col Nac Vicente Rocafuerte. 10: 32–35.

11. Campos FR (1930) Contribución del estudio de los mosquitos que habitan la ciudad y zonas adyacentes. Rev Col Nac Vicente Rocafuerte. 12: 40–41.

12. Campos FR (1930) Entomología Médica. Mosquitos peligrosos. Sava. Memoria referente al estudio de estos insectos. Rev Col Nac Vicente Rocafuerte. 12: 40–41.

13. Levi-Castillo R (1952) Redescription of *Aedes (Ochlerotatus) camposanus* Dyar (1918) as a valid species found in the coastal plain of Ecuador. Pacif Sci. 6:262–264.

14. Levi-Castillo R (1949) Atlas de los anofelinos Sudamericanos. Vol. 2. Sociedad Filantrópica del Guayas. Annals of the Entomological Society of America Press, Guayas.

15. Levi-Castillo R (1958) Provisional List of the Culicidae, Simuliidae, Phlebotomus and Culicoides of Ecuador. Proc Tenth Int Congr Entomol. 3.

16. Heinemann SJ, Belkin JN (1979) Collection records of the project “Mosquitoes of Middle America”. 13. South America: Brazil (BRA, BRAP, BRB), Ecuador (ECU), Peru (PER), Chile (CH). Mosq Syst. 11: 61–118.

17. GBIF-Secretariat (2020) Global Biodiversity Information Repository. Universitet sparken, Copenhagen, Denmark. Available at: https://www.gbif.org/es/.

18. Bisby FA, Roskov YR, Orrell TM, Nicolson D, Paglinawan LE, Bailly N, Kirk PM, Bourgoin T, Baillargeon G., eds (2010) Species 2000 & ITIS Catalogue of Life: 2010 Annual Checklist Taxonomic Classification. DVD; Species 2000: Reading, UK. Available at: http://www.catalogueoflife.org/annual-checklist/2010.

19. Roskov Y, Kunze T, Orrell T, Abucay L, Paglinawan L, Culham A, Bailly N, Kirk P, Bourgoin T, Baillargeon G, Decock W, De Wever A, Didžiulis V, eds (2014) Species 2000 & ITIS Catalogue of Life, 2014 Annual Checklist. Naturalis, Leiden, the Netherlands. Available at: https://www.catalogueoflife.org/annual-checklist/2014.

20. Harbach RE, Kitching IJ (1998) Phylogeny and classification of the Culicidae (Diptera). Syst Entomol. 23: 327–370.

21. Reinert JF (2001) Revised list of abbreviations for genera and subgenera of Culicidae (Diptera) and notes on generic and subgeneric changes. J Am Mosq Control Assoc. 17: 51–55.

22. Reinert JF (2009) List of abbreviations for currently valid generic-level taxa in family Culicidae (Diptera). Eur Mosq Bullettin 27: 68–76.

23. Harbach RE (2007) The Culicidae (Diptera): A review of taxonomy, classification and phylogeny. Zootaxa. 638: 591–638.

24. Kitching IJ, Lorna Culverwell C, Harbach RE (2015) The phylogenetic conundrum of *Lutzia* (Diptera: Culicidae: Culicini): a cautionary account of conflict and support. Insect Syst. Evol 46: 269–290.

25. Harbach RE (2013). The phylogeny and classification of Anopheles. In: Manguin S (Ed.), Anopheles mosquitoes- New insights into Malar Vectors. 3–53.

26. Obando RG, Carrejo NS (2009) Introducción al Estudio Taxonómico de Anopheles de Colombia. Programa Editorial Universidad del Valle Press, Colombia.

27. Rozo-Lopez P, Mengual X (2015) Updated list of the mosquitoes of Colombia (Diptera: Culicidae). Biodivers. data J.: e4567.

28. Calderón G, Fernández R, Valle J (1995) Especies de la fauna anofelina, su distribución y algunas consideraciones sobre su abundancia e infectividad en el Perú. Rev Peru Epidemiol. 8 (1): 5–23.

29. Simmons JS (1936) *Anopheles (Anopheles) punctimacula* naturally infected with malaria plasmodia. Amer Jour Trop Med. 16: 2–1.

30. Hayes J, Calderón G, Falcon R, Zambrano V (1987) Newly incriminated anopheline vectors of human malaria parasites in Junin Department, Peru. J Am Mosq Control Assoc. 3: 418–422.

31. Sinka ME (2013) Global Distribution of the Dominant Vector Species of Malaria, Anopheles mosquitoes - New insights into malaria vectors, Sylvie Manguin. IntechOpen. DOI: 10.5772/54163. Available from: https://www.intechopen.com/books/anopheles-mosquitoes-new-insights-into-malaria-vectors/global-distribution-of-the-dominant-vector-species-of-malaria

32. Conn JEL, Quiñones M, Marinete M (2013) Phylogeography, Vectors and Transmission in Latin America, Anopheles mosquitoes - New insights into malaria vectors, Sylvie Manguin. IntechOpen. DOI: 10.5772/55217.

33. Montoya-Lerma J, Solarte Y, Giraldo-Calderón GI, Quiñones ML, Ruiz-López F, Wilkerson RC, González R (2011) Malaria vector species in Colombia: A review. Mem Inst Oswaldo Cruz. 106: 223–238.

34. Zimmerman RH (1992) Ecology of malaria vectors in the Americas and future directions. Mem Inst Oswaldo Cruz. 87: 371–383.

35. Levi-Castillo R (1953) Lista provisional y distribución de los mosquitos Culicinos en el Ecuador. Rev Ecuat Entomol y Parasitol. 1: 34–45.

36. Ramírez N (1953) Especies Anofelinii predominantes en el Oriente Ecuatoriano. Oficina Sanitaria Panamericana Bolivia. 32(5): 499–504.

37. Calisher CH, Lazuick JS, Justines G, Francy DB, Monath TP, Gutierrez E, Sabattini MS, Bowen GS, Jakob WL (1981) Viruses isolated from *Aedeomyia squamipennis* mosquitoes collected in Panama, Ecuador, and Argentina: establishment of the Gamboa serogroup. Am J Trop Med Hyg. 30: 219–223.

38. Mitchell CJ, Monath TP, Sabattini MS, Cropp CB, Daffner JF, Calisher C, Jakob WL, and Christensen HA (1985) Arbovirus Investigations in Argentina, 1977–1980: II. Arthropod Collections and Virus Isolations from Argentine Mosquitoes. Am J Trop Med Hyg. 34: 945–955.

39. Gabaldon A, Ulloa G, Godoy N, Marquez E, Pulido J (1977) *Aedeomyia squamipennis* (Diptera, Culicidae) vector natural de malaria aviaria en Venezuela. BMSA. 17: 9–13.

40. Cevallos V, Ponce P, Waggoner JJ, Pinsky BA, Coloma J, Quiroga C, Morales D, Cárdenas MJ (2018) Zika and Chikungunya virus detection in naturally infected *Aedes aegypti* in Ecuador. Acta Trop. 177:74–80.

41. Myles, KM, Pierro DJ, Olson KE (2003) Deletions in the putative cell receptor-binding domain of Sindbis virus strain MRE16 E2 glycoprotein reduce midgut infectivity in *Aedes aegypti*. J Virol. 77: 8872–8881.

42. Pierro DJ, Myles KM, Foy BD, Beaty BJ, Olson KE (2003) Development of an orally infectious Sindbis virus transducing system that efficiently disseminates and expresses green fluorescent protein in *Aedes aegypti*. Insect Mol Biol. 12: 107–116.

43. Cancrini G, Di Regalbono AF, Ricci I, Tessarin C (2003) *Aedes albopictus* is a natural vector of *Dirofilaria immitis* in Italy. Vet Parasitol. 118:195–202.

44. Weaver SCR, Salas R, Rico-Hesse GV, Ludwig MS, Oberste J, Boshell J, Tesh RB (1996) VEE Study Group Re-emergence of epidemic Venezuelan equine encephalomyelitis in South America. Lancet. 1348(9025):436–40.

45. Rivas F, Diaz LA, Cardenas VM, Daza E, Bruzon L, Alcala A, De la Hoz O, Caceres FM, Aristizabal G, Martinez JW, et al (1997) Epidemic Venezuelan equine encephalitis in La Guajira, Colombia. J Infect Dis. 1175(4):828–32.

46. Eastwood G, Goodman SJ, Cunningham AA, Kramer LD (2013) *Aedes taeniorhynchus* vectorial capacity informs a pre-emptive assessment of West Nile virus establishment in Galápagos. Sci Rep. 3: 1–8.

47. Vasconcelos PFC, Sperb AF, Monteiro HAO, Torres MAN, Souza MRS, Vasconcelos HB, Mardini LH, Rodrigues SG (2003) Isolations of yellow fever virus from *Haemagogus leucocelaenus* in Rio Grande do Sul State, Brazil, in the Southern Cone. Trans R Soc Trop Med Hyg. 97.

48. Cardoso J, de Almeida MAB, dos Santos E, da Fonseca DF, Sallum MAM, Noll CA, de Monteiro HA, Cruz ACR, Carvalho VL, Pinto EV, et al (2010) Yellow Fever Virus in *Haemagogus leucocelaenus* and *Aedes serratus* mosquitoes, Southern Brazil. Emerg Infect Dis. 16:1918–1924.

49. Johnson BW, Cruz C, Felices V, Espinozoa WR, Mancok SR, Guevara C, Olson JG, Kochel TJ (2007) Ilheus virus isolate from a human, Ecuador. Emerg Infect Dis. 13: 956–958.

50. Whiteman NK, Goodman SJ, Sinclair BJ, Walsh T, Cunningham AA, Kramer LD, Parker PG (2005) Establishment of the avian disease vector *Culex quinquefasciatus* Say, 1823 (Diptera: Culicidae) on the Galápagos Islands, Ecuador. Ibis (Lond. 1859). 147: 844–847.

51. Consoli AGB, de Oliveira RL (1998) Principais mosquitos de importância sanitária no Brasil. Rio de Janeiro. Editora Fiocruz. 228 p.

52. Rutledge CR, Day JF, Lord C, Stark LM, Tabachnick WJ (2003) West Nile virus infection rates in *Culex nigripalpus* (Diptera: Culicidae) do not reflect transmission rates in Florida. J Med Entomol. 40:253–258.

53. Scherer WF, Dickerman RW, La Fiandra RP, Wong Chia C, Terrian J (1971) Infections of wild mammals: ecologic studies of Venezuelan encephalitis virus in southeastern Mexico. Am J Trop Med Hyg. 20: 980–988.

54. Evangelista J, Cruz C, Guevara C, Astete H, Carey C, Kochel TJ, Morrison AC, Williams M, Halsey ES, Forshey BM (2013) Characterization of a novel flavivirus isolated from *Culex* (Melanoconion) *ocossa* mosquitoes from Iquitos, Peru. J Gen Virol. 94: 1266–1272.

55. Weaver SC, Ferro C, Barrera R, Boshell J, Navarro JC (2004) Venezuelan Equine Encephalitis. Ann Rev Entomol. 49: 141–74.

56. Adames AJ (1971) XXIV A revision of the crabhole mosquitoes of the Genus Deinocerites. Contrib Am Entomol Inst. 7(2):1–15.

57. Njabo KY, Cornel AJ, Sehgal RN, et al (2009) *Coquillettidia* (Culicidae, Diptera) mosquitoes are natural vectors of avian malaria in Africa. Malar J. 8: 193.

58. Lebl K, Zittra C, Silbermayr K, Obwaller A, Berer D, Brugger K, Walter M, Pinior B, Fuehrer HP, Rubel F (2015) Mosquitoes (Diptera: Culicidae) and their relevance as disease vectors in the city of Vienna, Austria. Parasitol Rer. 114(2): 707–713.

59. Mahy BWJ, Van Regenmortel MHV (2008) Encyclopedia of virology (3rd ed. / editors-in-chief Brian W.J. Mahy and Marc H.V. Van Regenmortel). Academic Press, London.

60. Forattini OP (1965) Entomología Médica, Culicini: Culex, Aedes e Psorophora. Vol. 3. Universidade de São Paulo Press, São Paulo.

61. Lourenço-de-Oliveira R, Da Silva T (1985) Alguns aspectos da ecologia dos mosquitos (Diptera: Culicidae) de uma área de planicie (Granjas Calábria), em Jacarepaguá, Rio de Janeiro, III. Preferência horária das fêmeas para o hematofagismo. Mem Inst Oswaldo Cruz. 80(2):195–200.

62. Lourenço-de-Oliveira R, Heyden R (1986) Alguns aspectos da ecologia dos mosquitos (Diptera: Culicidae) de uma área de planicie (Granjas Calábria), em Jacarepaguá, Rio de Janeiro, IV. Preferência alimentares quanto ao hospedeiro e freqüência domiciliar. Mem Inst Oswaldo Cruz. 81(1):15–27.

63. Hervé J, Dég allier N, Travassos Da Rosa A, Pinheiro F, and Sá Filho G (1986) Arboviroses. Aspectos ecológicos. In: Instituto Evandro Chagas. 50 años de contribuição às Ciências Biológicas e à Medicina Tropical. Fund Serv Saúd Púb. 1:529–556.

64. Zavortink T (1979) The new sabethine genus *Johnbelkinia* and a preliminary reclassification of the composite genus *Trichoprosopon*. Contrib Am Entomol Inst. 17: p1–61.

65. Machado-Allison C, Barrera R, Delgado L, Gomez-Cova C, Navarro J (1986) Mosquitos (Diptera: Culicidae) de los Fitotelmata de Panaquire. Acta Biol Venez. 12: 1–12.

66. Barreto P, Lee VH (1969) Artrópodos Hematófagos del Río Raposo, Valle, Colombia: II - Culicidae. Caldasia. 49: 407–440.

67. De Rodaniche E, Galindo P (1957) Isolation of yellow fever virus from *Haemagogus mesodentatus, H. equinus* and *Sabethes chloropterus* captured in Guatemala in 1956. Am J Trop. Med Hyg. 6: 232–237.

68. Mittermeier RA, Myers N, Thomsen JB, et al (1998) Biodiversity hotspots and major tropical wilderness areas: approaches to setting conservation priorities. Conserv Biol. 12: 516–520.

69. Foley DH, Rueda LM, Wilkerson RC (2007) Insight into global mosquito biogeography from country species records. J Med Entomol. 44(4):554–567.

70. WHO (2015) World Malaria Report. Geneva: World Health Organization, Geneva, Suiza. Available at: https://apps.who.int/iris/bitstream/handle/10665/200018/9789241565158_eng.pdf;jsessionid=068D76F50FB992C5D54BCA8B8B9A9FEF?ua=1?sequence=1.

71. González R, Carrejo N, Wilkerson RC, Alarcon J, Alarcon-Ormasa J, Ruiz F, et al (2010) Confirmation of *Anopheles (Anopheles) calderoni* Wilkerson, 1991 (Diptera: Culicidae) in Colombia and Ecuador through molecular and morphological correlation with topotypic material. Mem Inst Oswaldo Cruz. 105: 1001–1009.

72. Pinault L, Hunter F (2011) New highland distribution records of multiple Anopheles species in the Ecuadorian Andes. Malaria Journal. 10: 236–246.

73. Gorham JR, Stojanovich CJ, Scott HG (1973) Clave ilustrada para los mosquitos anofelinos de Sudamérica occidental. Mosquito Systematics 5: 97–156.

74. Levi-Castillo R (1945) Anopheles pseudopunctipennis in the Los Chillos Valley of Ecuador, Journal of Economic Entomology. 38: 385–388.

75. Navarro JC, Arrivillaga J, Morales D, Ponce P, and Cevallos V (2015) Rapid assessment of mosquito biodiversity (Diptera: Culicidae) and health enviromental risk in a mountain area belongs to Ecuadorian chocó. Entomotropica. 30: 160–173.

76. Harrison BA, Ruiz-Lopez F, Calderon G (2012) *Anopheles (Kerteszia) lepidotus* (Diptera: Culicidae), not the malaria vector we thought it was: Revised male and female morphology; larva, pupa, and male genitalia characters; and molecular verification. Zootaxa. 3218: 1–17.

77. Forattini OP (1965) Entomología Médica. Vol. 1. Universidade de São Paulo Press, São Paulo.

78. Ruiz-Lopez F, Wilkerson R, Ponsonby D, et al (2013) Systematics of the Oswaldoi Complex (*Anopheles, Nyssorhynchus)* in South America. Parasites & Vectors. 6: 324.

79. Faran ME (1980) Mosquito Studies (Diptera: Culicidae) XXXIV.A revision of the Albimanus Section of the subgenus Nyssorhynchus of Anopheles. Contrib Am Entomol Inst Arbor. 155: 1–215.

80. Faran ME, Linthicum KJ (1981) A handbook of the Amazonian species of Anopheles (*Nyssorhynchus)* (Diptera: Culicidae). Mosq Syst. 13: 1–81.

81. Fritz G, Engman S, Rodriguez R, Wilkerson R (2004) Identification of four vectors of human *Plasmodium spp.* by multiplex PCR: *Anopheles rangeli, An. strodei, An. triannulatus, and An. trinkae* (Diptera: Culicidae: Nyssorhynchus). J Med Entomol. 41: 1111–1115.

82. Lounibos LP, Duzak D, Linley JR (1997) Comparative egg morphology of six species of the Albimanus section of Anopheles (Nyssorhynchus) (Diptera: Culicidae). J Med Ent. 34:136–55

83. Harbach RE, Howard TM (2009) Review of the genus Chagasia (Diptera: Culicidae: Anophelinae). Zootaxa. 2210: 1–25.

84. Sallum AM, Schultz T, Foster P, Aronstein K, Wirtz R, Wilkerson R (2002) Phylogeny of Anophelinae (Diptera: Culicidae) based on nuclear ribosomal and mitochondrial DNA sequences. Systematic Entomology. 27: 361–382.

85. Berlin OGW (1969) Mosquito studies (Diptera, Culicidae) XII. A revision of the neotropical subgenus Howardina of Aedes. Estudios sobre zancudos (Diptera, Culicidae) XII. Revisión del género neotropical Howardina de Aedes. Contrib Am Entomol Inst. 4: 1–190.

86. Navarro JC, Enríquez S, Arrivillaga J, Benitez Ortíz W (2016) Un nuevo *Aedes* para la Amazonía de Ecuador y actualización taxonómica del género para el país. Bol Malariol y Salud Ambient. 56: 113–121.

87. Campos FR (1922) Estudios sobre la Fauna Entomológica del Ecuador. 2° Dipteros Nematóceros: Fam. Culicidae (Mosquitos). Rev del Col Nac Vicente Rocafuerte. 1:18–30.

88. Arnell JH (1976) Mosquito studies (Diptera, Culicidae) XXXIII A revision of the Scapularis Group of Aedes (Ochlerotatus). Contrib Am Entomol Inst. 13: 1–144.

89. Bataille A, Cunningham AA, Cedeño V, et al (2009) Evidence for regular ongoing introductions of mosquito disease vectors into the Galapagos Islands. Proc Biol Sci. 276(1674): 3769–3775.

90. Bataille A (2012) Host selection and parasite infection in *Aedes taeniorhynchus*, endemic disease vector in the Galápagos Islands. Infection, Genetics and Evolution. 12: 1831–1841.

91. Siers S, Merkel J, Bataille A, Vargas FH, Parker PG (2010) Eccological correlates of microfilariae prevalence in endangered Galápagos birds. Journal of Parasitology. 96(2): 259–272.

92. Stone A, Knight KL, Starcke H (1959) A Synoptic Catalog of the Mosquitoes of the World (Diptera, Culicidae). Entomological Society of America, Press Thomas Say Foundation, Washington D.C.

93. Eastwood G, Kramer LD, Goodman SJ, Cunningham AA (2011) West Nile virus vector competency of *Culex quinquefasciatus* mosquitoes in the Galápagos Islands. Am J Trop Med Hyg. 85: 426–433.

94. Schick RX (1970) The Terrens group of Aedes (Finlaya). Contrib Amer Ent Inst. 5:1–158.

95. Levi-Castillo R (1954) Notes on Ecuadorian mosquitoes *Haemagogus equinus* in Ecuador and further taxonomic notes on *Uranotaenia aequatorianna* Levi-Castillo 1953. Rev Ecuat Entomol Parasitol. 2:83–86.

96. Arnell JH (1973) Mosquito studies (Diptera: Culicidae) XXXII. A revision of the genus Haemagogus. Contrib Amer Entomol Inst. 10:1–174.

97. Valencia JD (1973) Mosquito studies (Diptera, Culicidae) XXXI. A revision of the subgenus Carrolia of Culex. Contribution of the American Entomological Institute. 9 (4): 1–134.

98. Sirivanakarn S, Galindo P (1980) *Culex (Meknoconion) aukmesi.* a new species from Panama (Diptera: Culicidae). Mosq syst. 12:25–34.

99. Pecor JE, Mallampalli VL, Harbach RE, Peyton EL (1992) Catalog and illustrated review of the subgenus Melanoconion of *Culex* (Diptera: Culicidae). Contributions of the American Entomological Institute. 27(2): v1–228.

100. Forattini OP, Sallum MAA (1992) A new species of Culex (Melanoconion) from the amazonian region (Diptera: Culicidae). Mem Inst Oswaldo Cruz. 87: 265–274.

101. Sirivanakarn S, Belkin JN (1980) The identity of *Culex (Melanoconion) taeniopus* Dyar and Knab and related species with notes on the synonymy and description of a new species (Diptera: Culicidae). Mosq Syst. 12: 7–24.

102. Adames AJ, Arzube ME (1975) Geographical extension of *Galindomyia Zeei* Stone and Barreto to Ecuador (Diptera: Cuhcidae). Mosq Syst. 7:113–114.

103. Belkin J (1968) The type specimens of New World mosquitoes in European museums. Contrib Am Entomol Inst. 3: 1–72.

